# The Environmental, Endocrine, and Epigenetic Basis of Size Plasticity in a Superorganism

**DOI:** 10.64898/2026.09.17.752427

**Authors:** Navid Bahramifarid, Akash Dahiya, Vie Tran, Kami Fabien, Rajendhran Rajakumar

## Abstract

From plants to animals, genomes are responsive to environmental variation, the interaction of which can influence developmental processes and facilitate profound consequences on life-history and evolution. The interplay between genes and the environment is known to be mediated by the action of endocrine and epigenetic mechanisms including histone modifications. Ant development is plastic: environmental changes can induce diverse phenotypes, generating different castes that distinguish individuals by size, behaviour, and reproduction. While recent work has demonstrated that hormonal and epigenetic mechanisms underlie caste-specific trait variation in ants, the interplay of these mechanisms, and the process by which they mediate environmental changes to produce distinct trait variation, remains enigmatic. Here, we use the invasive fire ant *Solenopsis invicta*, dubbed a superorganism and known for its extreme worker caste size variation, to investigate the interplay between endocrine and epigenetic mechanisms in response to environmental variation. We found that sizing and developmental timing are both thermally and nutritionally plastic, influenced by juvenile hormone, ecdysone, histone (de)acetylation, and histone (de)methylation, and that these mediators interact and are thermally and nutritionally plastic. Surprisingly, fire ants break the Temperature-Size Rule, ecdysone makes ants bigger faster, and histone methylation disruption unexpectedly makes ants smaller. These and other findings suggest that an endocrine-epigenetic axis mediates thermal and nutritional plasticity underlying the extreme size variation that has evolved in the complex worker caste system in fire ants. More generally, we propose that an environment-endocrine-epigenetic (E3) approach will provide an integrative perspective on the development and evolution of adaptive phenotypes.

## INTRODUCTION

Whether nutritional variation or climate change, organisms need to adapt to a changing environment^1^. The ability of an organism to developmentally respond to environmental variation generating diverse phenotypes is called developmental plasticity^1–3^. On a molecular level, plants and animals can plastically respond to environmental variation through endocrinological and epigenetic mechanisms^1,4,5^. Environmental variation can include a wide array of both abiotic cues such as drought and temperature, and biotic cues such as nutrition and predators^6^. For example, in the red-eared slider turtle *Trachemys scripta*, sex is determined by temperature variation, and this is regulated both hormonally through the sex steroid hormones (estrogens and androgens), and epigenetically through the histone modifier lysine demethylase 6b (*Kdm6b*)^7–9^. Furthermore, in the honeybee *Apis mellifera,* caste (queen versus worker) is determined by nutritional variation, and this is regulated both hormonally through juvenile hormone (JH), and epigenetically through DNA methylation and microRNAs (miRNAs)^10–13^. A holistic approach espoused by Lewontin in the Triple Helix^14^, which factors in the environment, gene activity during development, and phenotypic variation is needed to understand the origins of biological diversity. Based on this, finding a model organism that integrates studies of environmental variation, and both endocrinological and epigenetic regulation (E3) of phenotypic variation would provide fundamental insight into how organisms developmentally respond and evolutionarily adapt to their environment.

*Solenopsis invicta* is a model ant species, both in basic research and in applied pest-eradication contexts providing an ideal starting point for an E3 approach. *S. invicta* is a highly invasive species native to South America that is estimated to have first invaded North America in the early 20^th^ century and are in the top 10 list of most economically damaging species recorded, at a total cost of $16.7 billion USD^15,16^. Fire ants live in colonies that has a reproductive queen caste and a sterile worker caste that varies dramatically in their sizing from very small workers to massive-sized workers in the form of a size continuum (Fig.1*A*)^15,17^. Akin to the differentiation of germ cells and specialized somatic cells in an organism, fire ant colonies composed of specialized reproductive and sterile castes, which interact and coordinate their tasks, are thought of as a superorganism^18,19^. In ants, rather than a genetic basis, this phenotypic variation is due to developmental plasticity^20,21^. Caste-specific phenotypic variation is generated through the action of environmental variation on developmental endocrinological and epigenetic processes^22–27^.

**Fig. 1.**
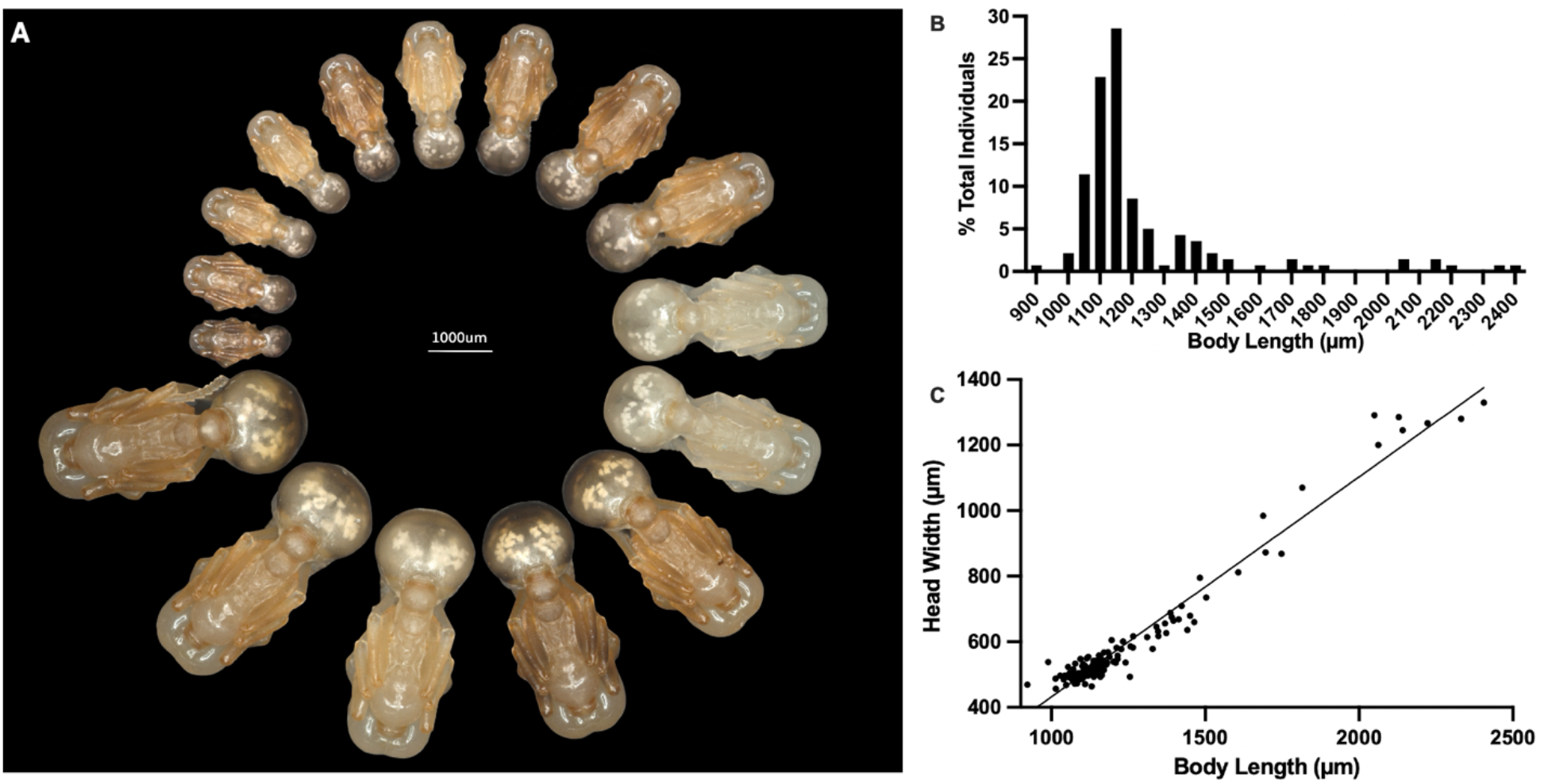
*Solenopsis invicta* natural size and allometry across the worker size continuum. **(A)** Pupal size pinwheel representing the general size continuum of differently sized individuals at their pupal stage within the *S. invicta* worker caste, separated by intervals of ∼100 um within a fire ant colony. **(B)** Frequency distribution diagram showing % of body sizes across individuals. **(C)** An XY scatterplot to compare the head width-to-body length distribution of individuals sampled from a fire ant colony.

This size diversity found in *S. invicta* is critical for their colony’s efficiency in the division of labour as different-sized individuals are adapted to, and perform, specialized tasks to varying frequencies^17,28,29^. Across fire ant development, an individual progresses from an embryo to four separate larval instars until they metamorphose into a pupa and subsequently eclose into an adult ant^15,30^. During embryogenesis, optimal temperature and photoperiod induce embryos to become queens, while later during larval development, individuals are thought to undergo varying duration/timing of their last instar to enable the generation of differently sized workers^15,31^. As ants are a holometabolous insect, this final instar growth determines the final size of the adult individual, which is reflected in the pupae after metamorphosis (Fig1*A-C*)^26,31,32^.

To better understand the mechanisms that underlie developmental plasticity in *S. invicta* that have facilitated their ecological and invasive success, we have taken an E3 approach exploring how variations in environmental factors and their underlying endocrine and epigenetic mediators influence phenotypic variation. To do this, we used sizing (head size, body size, and allometry) and developmental timing as phenotypic readouts following manipulations in temperature, nutrition, JH and ecdysone hormone, and histone acetylation and methylation. Furthermore, we investigated the influence of temperature and nutrition on hormonal signaling and histone modifier activity, and the interplay between hormones and epigenetic modifiers. By taking an E3 approach, we were able to identify unexpected roles of the **E**nvironment (temperature), as well as **E**ndocrine (ecdysone) and **E**pigenetic (histone methylation) factors involved in generating the size continuum of *S. invicta* workers. Altogether, an integrative E3 approach has illuminated how endocrine and epigenetic mechanisms connect fire ants to their changing environment.

## RESULTS

### Thermal Plasticity and the Epi-Endocrine Axis in *S. invicta*

Variation in thermal conditions is known to affect development in *S. invicta* colonies^33^. Specifically, higher temperatures led to exponentially more rapid growth of developing brood (embryos, larvae, pupae), with an optimum of colony size at 32°C. To better understand the role of thermal plasticity on the worker development we characterized the effects of temperature variation on size and developmental timing, as well as potential endocrine and epigenetic modifiers underlying this plasticity. We reared *S. invicta* early 4^th^ instar larvae at low (25°C), medium (28.5°C), and high (32°C) temperatures, as this is the developmental period when the majority of growth initiates, and characterized head width and body size of pupae. Surprisingly, relative to the 25°C group, we found increases for 28.5°C reared individuals for both head width (Fig. 2*A*) and body length (Fig. 2*B*). Next, we compared the differences in developmental timing between larvae reared among the three temperatures and found that increasing temperatures shortens the total developmental period (Fig. 2*C*). Finally, to assess the influence of temperature on head-to-body-allometry, we compared head size to body size at the different temperatures. While temperature did not influence the head-to-body allometric slope across temperatures, we found that a decrease between the intercepts of the 28.5°C treatment, relative to the 25°C and 32°C groups (Fig. 2*D*). Across holometabolous insects, increased temperature causing a shortening of developmental timing and a reduction in organismal size is nearly universal and known as the temperature size rule (TSR)^34^. Taken together, temperature increase is unexpectedly associated with an increase in head and body size as well as rapid development.

**Fig. 2.**
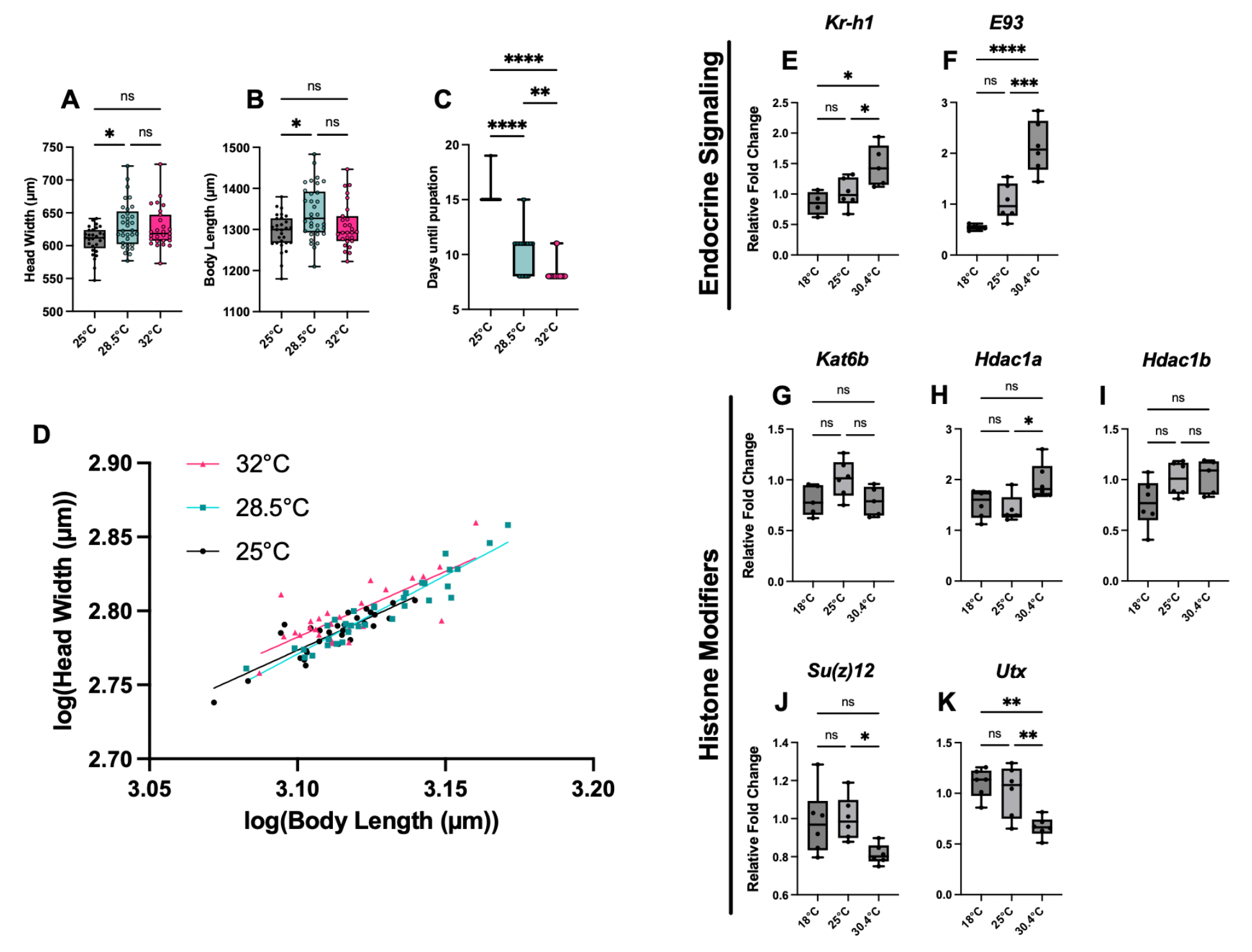
The effects of temperature variation on developmental plasticity in *Solenopsis invicta*. Comparison between measurements of **(A)** head width, **(B)** body length, **(C)** developmental timing, and **(D)** head-to-body allometry of developing *S. invicta* larvae in 25°C (n=29), 28.5°C (n=33), and 32°C (n=27) treatment groups. qPCR analyses of 48-hour treatments comparing **(E)** *Kr-h1*, **(F)** *E93*, **(G)** *Kat6b*, **(H)** *Hdac1a*, **(I)** *Hdac1b*, **(J)** *Su(z)12*, and **(K)** *Utx* between 18, 25, and 30.4°C groups. Bars represent means, with standard deviation (SD) indicated with error bars. Values for statistical significance of ANOVA and Kruskal-Wallis tests include \**p*<0.05, \*\**p*<0.01, \*\*\**p*<0.001, and \*\*\*\**p*<0.0001. Sample size for qPCR treatment groups range from 4-6 biological replicates, each containing 5 individual larvae.

Based on our findings of thermal plasticity of size and developmental timing, we next wanted to characterize the effects of temperature on both the endocrine and epigenetic axes underlying fire ant size variation. In holometabolous insects, juvenile hormone (JH) and ecdysone are the master ancient regulators of developmental timing, growth, and metamorphosis in insects^35,36^. Therefore, in the context of the endocrine axis, we measured the expression of *Kr-h1*, the key effector of JH signalling pathway, and *E93*, a critical effector of the ecdysone signalling pathway^37^. To do this, early 4^th^ instar larvae were reared at low (18°C), medium (25°C), and high (30.4°C) temperatures, and then expression levels of the endocrine (*Kr-h1*, *E93*) were measured. We found that increasing temperatures resulted in the increased expression of both *Kr-h1* (Fig. 2*E*) and *E93* (Fig. 2*F*). Therefore, there is a positive influence of temperature on the two essential developmental hormones in the fire ants.

In the context of the epigenetic axis, we measured the expression of histone epigenetic toolkit genes, specifically the acetyltransferase *Kat6b* (also known as *enok*) and the histone deacetylase paralogs *Hdac1a* and *Hdac1b*, essential writers/erasers of histone acetylation as well as histone methyltransferase *Su(z)12* and histone demethylase *Utx*, essential writers/erasers of histone methylation^38–41^. We selected these histone modifiers as they have been implicated both in the ant comparative genomic literature as candidate genes facilitating adaptive radiations, size variation, and caste diversity^42,43^, and the ant transcriptomic literature as candidate mediators of environmental response and caste plasticity^44–46^. Furthermore, these histone modifiers exhibit dynamic expression levels across key stages of fire ant worker larval development (Fig. S1*A-E*). For histone acetylation modifiers, *Kat6b* did not respond to temperature variation (Fig. 2*G*), and while expression levels of *Hdac1a* increased with temperature (Fig. 2*H*), *Hdac1b* was also unaffected (Fig. 2*I*). This divergence in transcriptional response of these *Hdac1* paralogs is consistent with our observations that they have different expression dynamics during the critical stages of larval development (Fig. S1*B, C*). For histone methylation modifiers, both *Su(z)12* and *Utx* decreased with temperature (Fig. 2*J*, *K*). Collectively, at the phenotypic level, higher temperatures can increase sizing of individuals and shorten developmental timing, while at the molecular level, ecdysone signalling as well as histone deacetylation modifiers increase with temperature while histone (de)methylation modifiers decrease.

### Nutritional Plasticity and the Epi-Endocrine Axis in *S. invicta*

Nutrition is known to influence queen-worker and worker-soldier caste development its allocation across the colony is tightly regulated in ants^22,23,47–49^. Following our E3 approach with fire ants, we therefore looked at nutrition to investigate the influence of nutritional plasticity on growth, developmental timing, as well as the endocrine and epigenetic axes. We first reared *S. invicta* early 4^th^ instar larvae in either an intermittently fasted group with or a fed group and characterized head width and body size, developmental timing, and head-to-body allometry of pupae. Head, body size, and allometry remained unchanged in the fasting group as compared to the sated group (Fig. 3*A*, *B*, *D*). Despite the lack of change in sizing, intermittent fasting individuals prolonged development compared to the fed group (Fig. 3*C*). This developmental timing extension likely is the cause for intermittently fasted individuals catching up in size to the fed group. To further investigate nutrition-dependent size variation, we tried an alternative nutritional regime of nutrient deprivation, where individuals were fed only sugar water and compared to fed individuals. While this nutritional regime led to a high rate of mortality, four individuals survived, and both their head and body sizes were dramatically smaller than the fed group, such that they were smaller than any individuals found in the natural size distribution (Fig. 1*A*-*C*; Fig. S*2*). Furthermore, this change in size was mirrored by a dramatic increase in developmental timing, prolonging development of nutrient-deprived individuals by an extra 2-5 weeks as compared to fed controls (2-to-3-fold extension). Based on our findings of nutritional plasticity of size and developmental timing, we reared early 4^th^ instar larvae in either a fed or starvation condition and then assessed the expression of the endocrine and histone modifier toolkit genes after a 96hr treatment. For endocrine signalling, while starvation did not lead to a change in *Kr-h1* expression relative to fed controls (Fig. 3*E*), *E93* decreased in expression levels (Fig. 3*F*). For histone acetylation modifiers, while *Kat6b* did not change following starvation (Fig. 3*G*), both *Hdac1a* (Fig. 3*H*) and *Hdac1b* (Fig. 3*I*) decreased in expression levels compared to fed controls. For histone methylation modifiers, *Utx* decreased in expression levels following starvation, as compared to fed controls (Fig. 4*K*), while *Su(z)12* did not change (Fig. 3*J*). Altogether, at the phenotypic level, nutritional deficiency prolongs developmental timing and can negatively affect growth, while at the molecular level, ecdysone signalling, as well as both histone acetylation and histone demethylation modifiers all decrease.

**Fig. 3.**
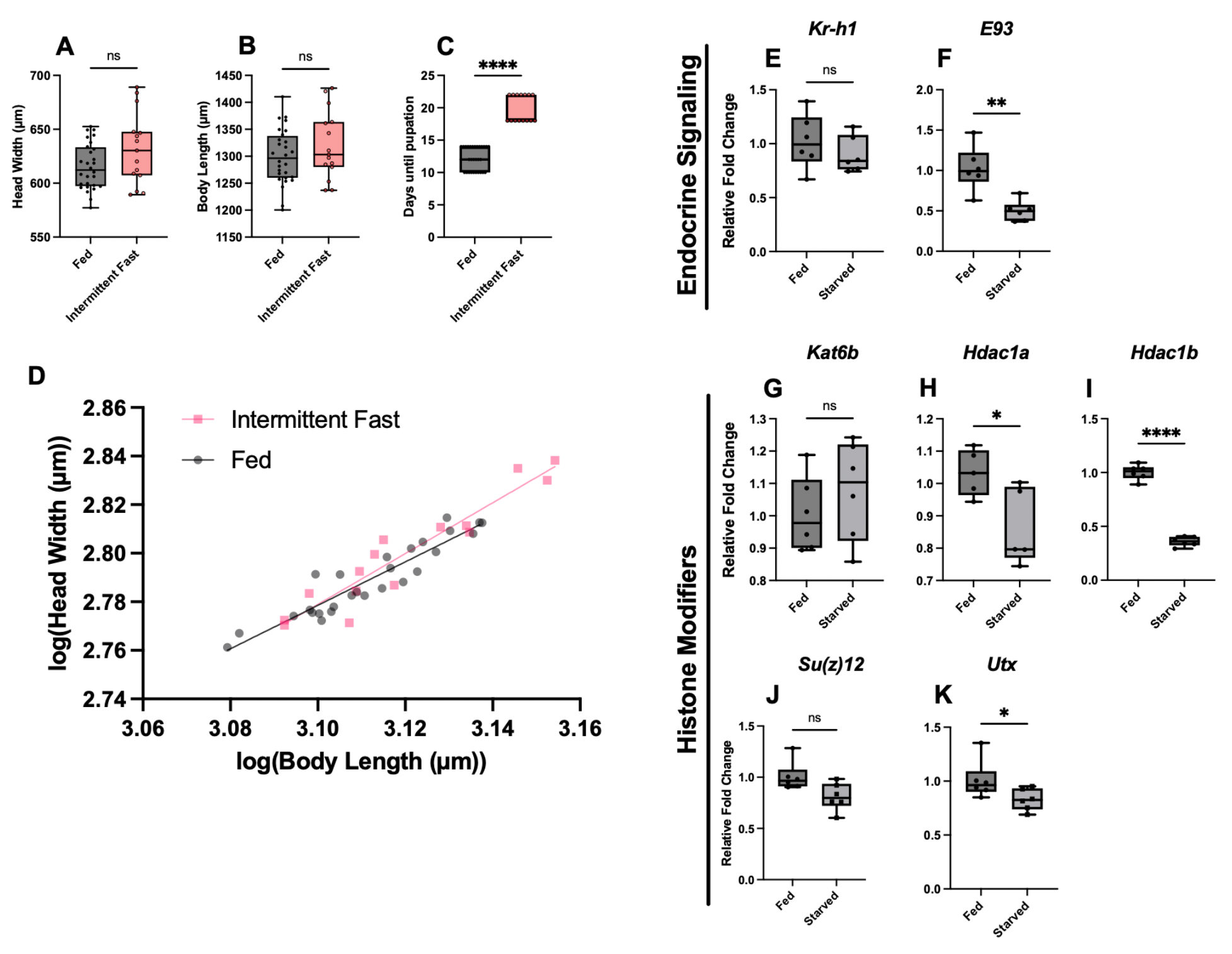
The effects of nutritional variation on developmental plasticity in *Solenopsis invicta*. Comparison between measurements of **(A)** head width, **(B)** body length, **(C)** developmental timing, and **(D)** head-to-body allometry of developing *S. invicta* larvae in sated (n=28) and intermittently fasted (n=15) treatment groups. qPCR analyses of 96-hour treatments comparing **(E)** *Kr-h1*, **(F)** *E93*, **(G)** *Kat6b*, **(H)** *Hdac1a*, **(I)** *Hdac1b*, **(J)** *Su(z)12*, and **(K)** *Utx* between sated and starved groups. Bars represent means, with standard deviation (SD) indicated with error bars. Values for statistical significance of Student’s *t*-tests and Mann-Whitney tests include \**p*<0.05, \*\**p*<0.01, and \*\*\*\**p*<0.0001. Sample size for qPCR treatment groups range from 5-6 biological replicates, each containing 5 individual larvae.

**Fig. 4.**
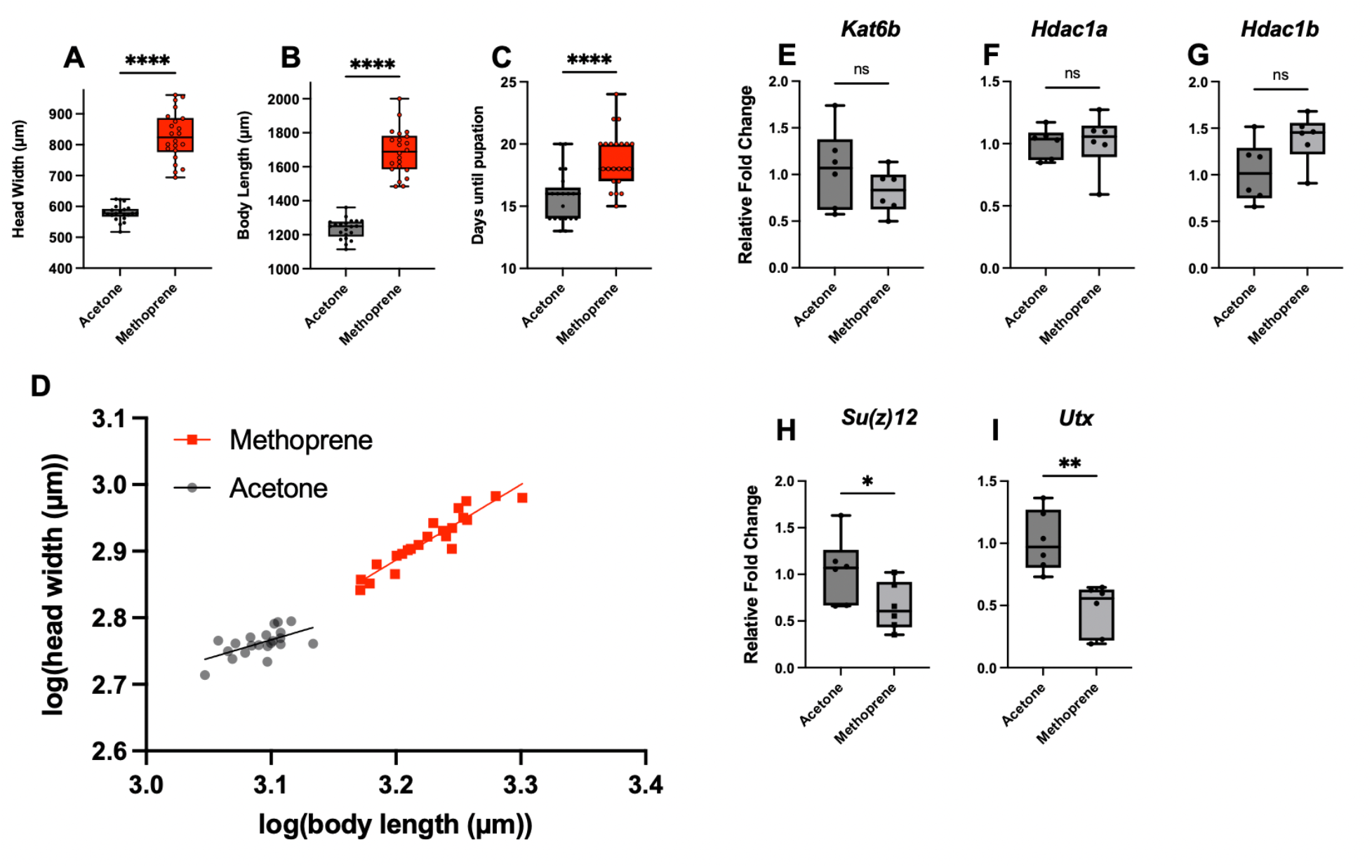
The effects of the juvenile hormone analogue methoprene on developmental plasticity in *Solenopsis invicta*. Comparison between measurements of **(A)** head width, **(B)** body length, **(C)** developmental timing, and **(D)** head-to-body allometry of developing *S. invicta* larvae in acetone controls (n=21) and methoprene (n=23) treatment groups. qPCR analyses of 24-hour treatments comparing **(E)** *Kat6b*, **(F)** *Hdac1a*, **(G)** *Hdac1b*, **(H)** *Su(z)12*, and **(I)** *Utx* between acetone and methoprene groups. Bars represent means, with standard deviation (SD) indicated with error bars. Values for statistical significance of Student’s *t*-tests include \*\**p*<0.01 and \*\*\*\**p*<0.0001. Sample size for qPCR treatment groups include 6 biological replicates, each containing 5 individual larvae.

### Endocrine Axis Influences Growth and the Epigenetic Axis in *S. invicta*

Juvenile hormone is known to influence size and developmental timing in *S. invicta* in a dose-dependent manner. Wheeler^31^ demonstrated that treatment of JH to developing *S. invicta* larvae led to an increase in body size and an extension of larval development. How JH influences epigenetic processes, and specifically histone modifiers is unknown. Therefore, we treated developing *S. invicta* individuals with JH analog (JHA) and assayed the influence of JH on sizing, head-to-body allometry, and histone modifier expression levels. We injected early 4^th^ instar larvae with JHA and as expected, we found that compared to the control, JHA increased head width and body size, as well as extended developmental timing (Fig. 4*A*, *B*, *C*). As the influence of JH on head-to-body allometry has yet to be described in fire ants, we therefore analyzed the allometric effects of JHA and found that treatments led to an increase in slope as compared to controls (Fig. 4*D*). Based on our findings of the effects of JH signalling on sizing, allometry, and developmental timing, we next treated early 4^th^ instar larvae with JHA and characterized the influence of JH signalling on the expression levels of histone modifiers. For histone acetylation modifiers, individuals treated with JHA for 24 hours had no effect on expression levels as compared to controls (Fig. 4*E-G*), while JHA treatment led to decreased expression of both histone methylation modifiers *Su(z)12* (Fig. 4*H*) and *Utx* (Fig. 4*I*). Altogether, at the phenotypic level, JH signalling prolongs developmental timing, positively affects sizing and allometry, while at the molecular level, histone acetylation modifiers are unresponsive to JH, yet histone methylation modifiers decrease.

The steroid hormone ecdysone, conserved across arthropods in regulating ecdysis, is the critical regulator of larval-to-larval moults, the timing for the cessation of growth, and the induction of metamorphosis in insects^36,50,51^. In ants, while very little has been done to assess the role of ecdysone during development, ecdysone has been shown in ants, to be negatively associated with queen development and a cause a decrease in both size and timing of developing queens^52–55^, consistent with ecdysone’s influence on developmental timing and growth in other insects^36,56–58^. To test the influence of ecdysone on fire ant sizing, developmental timing, allometry, and histone modifier activity, we injected early 4^th^ instar larvae with 20-hydroxyecdysone (20E), the active form of ecdysone. Consistent with that found in other insects, 20E shortened developmental timing (Fig. 5*C*). Surprisingly, rather than decreasing size, we found that, compared to controls, 20E increased both head (Fig. 5*A*) and body size (Fig. 5*B*). This change in size was proportional, as there were no changes in head-to-body allometry (Fig. 5*D*). Based on our findings of the effects of ecdysone signalling on sizing, allometry, and developmental timing, we next treated 20E to early 4^th^ instar larvae and described the influence of ecdysone on histone modifier expression levels. After a 24 hours treatment, larvae treated with 20E did not change in the activity of histone acetylation nor histone methylation modifier levels, as compared to controls (Fig. 5*E*-*I*). Therefore, the action of ecdysone signalling on sizing and developmental timing may be independent of histone modifiers. Altogether, we found that 20E treatments led to an unexpected increase in head and body sizes, and further, this increase in sizing occurred despite a shortening of developmental timing.

**Fig. 5.**
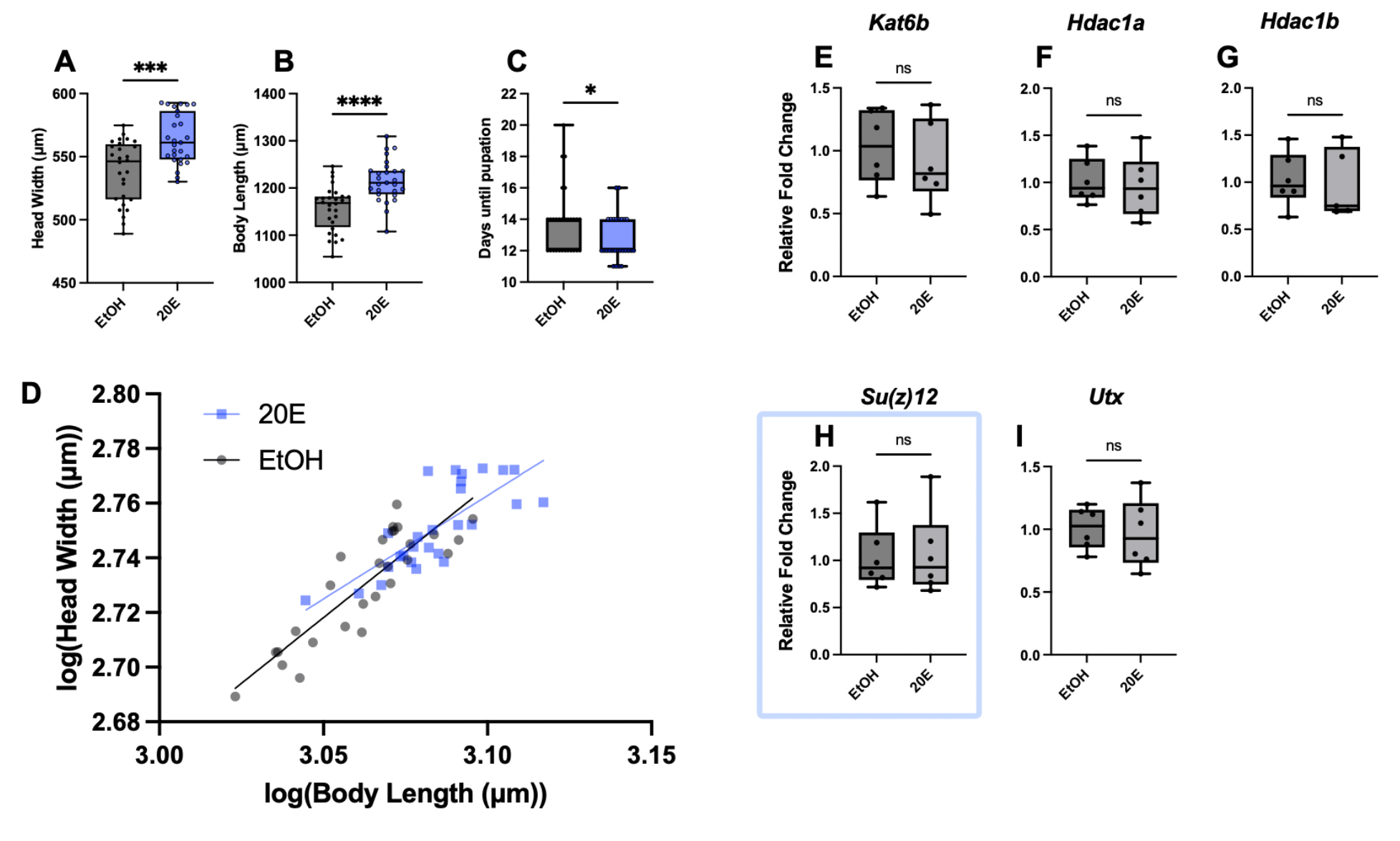
The effects of 20-hydroxyecdysone (20E) on developmental plasticity in *Solenopsis invicta*. Comparison between measurements of **(A)** head width, **(B)** body length, **(C)** developmental timing, and **(D)** head-to-body allometry of developing *S. invicta* larvae in ethanol controls (n=28) and 20E (n=27) treatment groups. qPCR analyses of 24-hour treatments comparing **(E)** *Kat6b*, **(F)** *Hdac1a*, **(G)** *Hdac1b*, **(H)** *Su(z)12*, and **(I)** *Utx* between ethanol and 20E groups. Bars represent means, with standard deviation (SD) indicated with error bars. Values for statistical significance of Student’s *t*-tests and Mann-Whitney test include \**p*<0.05, \*\*\**p*<0.001, and \*\*\*\**p*<0.0001. Sample size for qPCR treatment groups range from 5-6 biological replicates, each containing 5 individual larvae.

### Epigenetic Axis Influences Growth and the Endocrine Axis in *S. invicta*

To date, very little is known of the role of histone modifiers during ant development. Our findings have demonstrated that histone (de)acetylation and histone (de)methylation are responsive in varying degrees to environmental and hormonal variation and are dynamic in expression during caste-specific development. We therefore wanted to functionally manipulate this modification to investigate their role in sizing, allometry, and developmental timing. In parallel to the erasure activity of histone deacetylation, the writer activity of histone methylation coordinates with histone deacetylation, and typically they promote heterochromatin formation and transcriptional repression^59^. Therefore, to characterize the role of these heterochromatin-forming histone modifications, we used a pharmacological approach to inhibit them. Developing early 4^th^ instar larvae that were injected with the extensively used histone deacetylase inhibitor trichostatin A (HDACi, TSA) that has been previously tested in mammals, nematodes, flies, ants, and beetles^46,60–64^ led to an increase in head width (Fig. 6*A*) and body size (Fig. 6*B*), without altering developmental timing nor allometry (Fig. 6*C*, *D*), as compared to controls. This indicates that histone acetylation, known to regulate the activation of transcription of developmental genes through euchromatin formation^38,59,65^ is associated with a proportional increase in size. However, as shown above, starvation, negatively regulate size, yet is associated with a downregulation of histone deacetylases *Hdac1a* and *Hdac1b* (Fig. 3H, I). One explanation for this discrepancy is that both HDACi and starvation are known to induce autophagy, a cell survival process critical for the allocation of limited cellular resources and our HDACi was done on well-nourished individuals^66–68^. To see if HDACi individuals were benefitting from both nutrition and autophagy towards their size increase we measured the expression of the autophagy gene *atg8* which is known in plants and animals to transcriptionally respond to starvation conditions^69–71^. Comparing HDACi to controls, we found that *atg8* increased in expression (Fig. 6*E*).

**Fig. 6.**
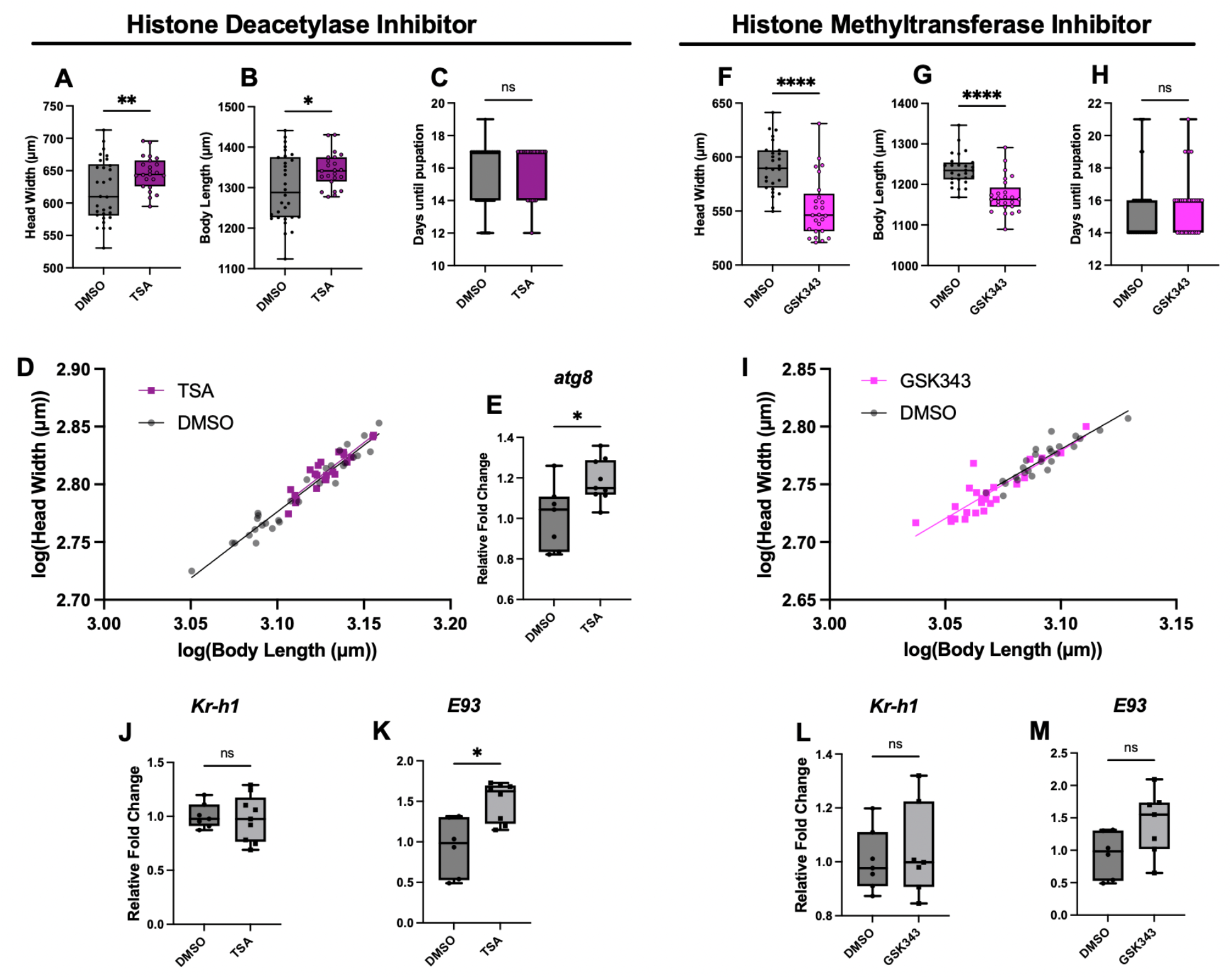
The effects of pharmacological inhibitors of histone modifications on developmental plasticity in *Solenopsis invicta*. Comparison between measurements of (A) head width, (B) body length, (C) developmental timing, and (D) head-to-body allometry of developing *S. invicta* larvae in DMSO controls (n=16) and histone deacetylase inhibitor TSA (n=13) treatment groups. qPCR analysis of a 48-hour treatment comparing (E) *atg8* between DMSO and TSA groups. Comparison between measurements of (F) head width, (G) body length, (H) developmental timing, and (I) head-to-body allometry of developing *S. invicta* larvae in DMSO controls (n=27) and histone methyltransferase inhibitor GSK343 (n=25) treatment groups. qPCR analyses of 48-hour treatments comparing (J) *Kr-h1* and (K) *E93* between DMSO and TSA groups, as well as (L) *Kr-h1* and (M) *E93* between DMSO and GSK343 groups. Bars represent means, with standard deviation (SD) indicated with error bars. Values for statistical significance of Student’s *t*-tests, Mann-Whitney tests, and Welch’s t tests include \**p*<0.05, \*\**p*<0.01 and \*\*\*\**p*<0.0001. Sample size for qPCR treatment groups range from 6-9 biological replicates, each containing 3 individual larvae.

Histone methylation can inactivate developmental gene transcription through the PRC2 complex and H3K27me3^38,72^. We therefore used a histone methyltransferease inhibitor (HMTi; GSK343), previously tested in nematodes and mammals^64,73^, to perturb this complex to see if it is a negative regulator of size in fire ants. Surprisingly, perturbing early 4^th^ instar larvae with the HMTi led to a decrease in size, HMTi-treated individuals were smaller in head width (Fig. 6*F*) and body sizes (Fig. 6*G*), without changing developmental timing (Fig. 6*H*) nor allometry (Fig. 6*I*), relative to controls. This indicates that histone methylation in fire ants can contribute towards positively regulating body size. Together, histone (de)acetylation and histone (de)methylation are histone epigenetic mechanisms that can influence size independent of developmental timing.

While our earlier findings suggest that developmental hormones can, to varying degrees, influence histone modifiers, very little is known of the capacity for histone modifiers to influence hormonal signalling in developing ants. Furthermore, given our findings of the effects of inhibitors of histone deacetylation and methylation modifiers on sizing, we next treated early 4^th^ instar larvae to determine the influence of HDACi and HMTi on the expression levels of hormonal signaling. After a 48-hour treatment, TSA treatments led to an increase in expression of *E93* (Fig. 6*K*), without influencing *Kr-h1* levels (Fig. 6*J*). Treatments with GSK343 did not affect expression levels of *Kr-h1* nor *E93* (Fig. 6*L, M*). Altogether, histone (de)acetylation and (de)methylation influence head and body size in a bidirectional manner, and histone deacetylation is negatively associated with an ecdysone activity.

## DISCUSSION

### Environmental Variation & Worker Polyphenism in Fire ants

#### Thermal Plasticity of Size and Developmental Timing

Navigating the E3 approach, when looking at E1 (Environment) in fire ants, we found that temperature increase led to an increase in size and a decrease in total duration of development (Fig. 2; Fig. 7, arrow 1). This finding is consistent with what has previously been demonstrated for both ant species *Pheidole pallidula* and *Camponotus floridanus*, where increasing temperature leads to an increase in growth and induction of the large soldier caste^23,74^. Complementarily, it has been shown that the incidence of high levels of worker size polymorphism is much higher in warm regions including tropical savannahs and the desert^75^. Together, these findings indicate that ants violate the Temperature Size Rule (TSR). Across holometabolous insects, TSR predicts that an increase in temperature would lead to a shorter duration of development, producing smaller individuals^34^. While fire ants are consistent with TSR when predicting temperature increase leading to a shorter period of development, they exhibit an unexpected increase in size. We propose that ants have co-opted aspects of hormonal signalling in such a way that increasing temperatures shortens developmental timing (consistent with TSR) and increases sizing (violates TSR) through an increased growth rate.

**Fig. 7.**
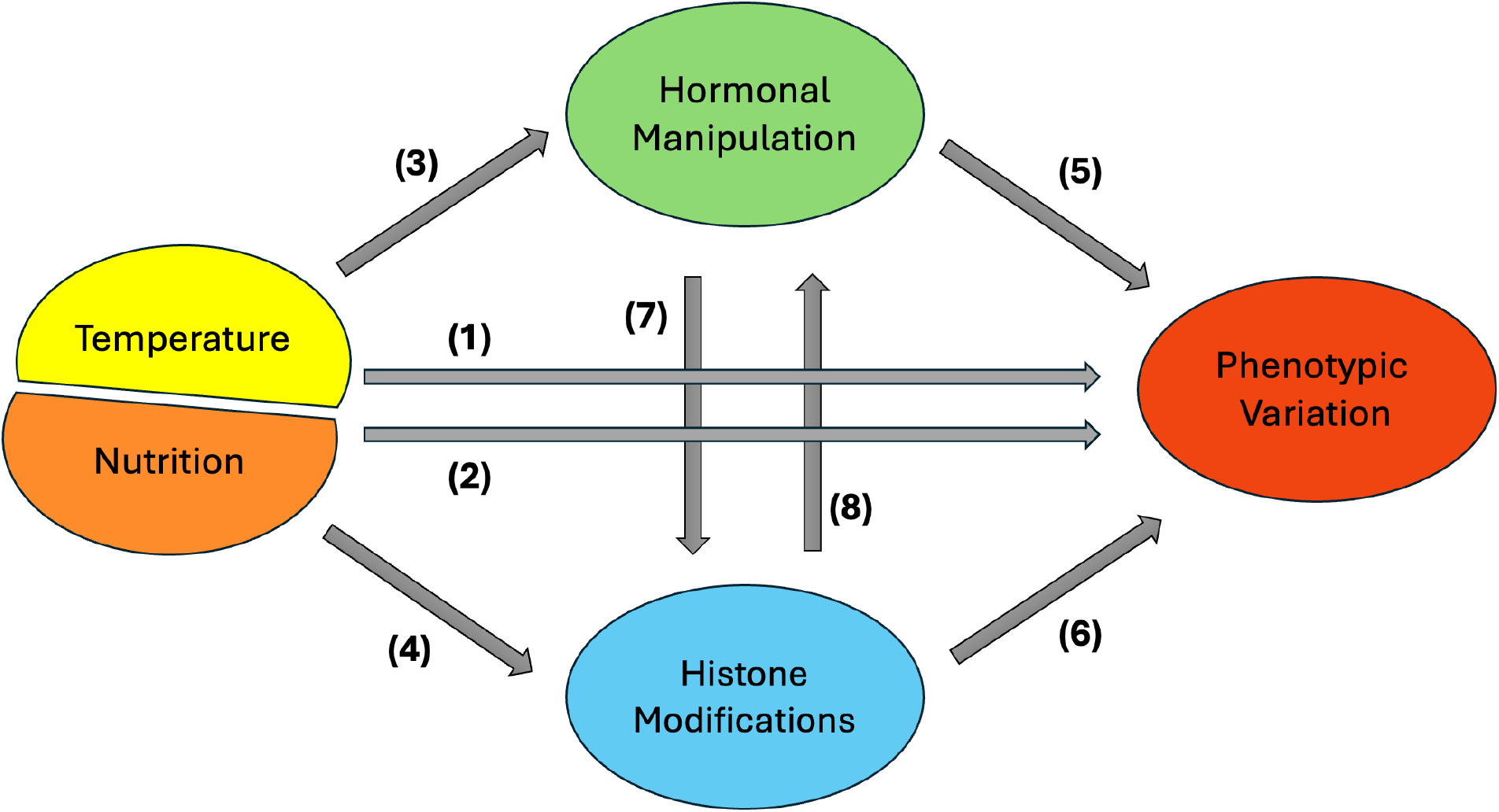
Connecting the environmental, endocrine, and epigenetic facets (E3) of developmental plasticity in *Solenopsis invicta*. Each facet is represented by their respective colours and are linked by arrows to indicate their interactions within fire ants. Full lines demonstrate relationships that have been characterized across *S. invicta* development. Each number represents the interplay between each respective facet. (1) Effects of changes in temperature on phenotypic variation. (2) Effects of changes in nutrition on phenotypic variation. (3) Effects of changes in temperature and nutrition on hormonal signalling. (4) Effects of changes in temperature and nutrition on histone modifier activity. (5) Effects of hormonal manipulation on phenotypic variation. (6) Effects of histone modifications on phenotypic variation. (7) Effects of hormonal manipulations on histone modifier activity. (8) Effects of histone modifications on hormonal signalling.

In mosquitoes, there is an exception to the TSR rule where individuals are smaller at higher temperatures when food quality is low, in contrast, when food is abundant, individuals grow larger at higher temperatures^76^. Fire ants can thermoregulate at the colony level in their mounds by cycling position of brood along a temperature gradient across the depth of the mound^77^. Along the thermal gradient of the colony, developing fire ant larvae experience parental care and feeding by the adult workers, potentially mirroring the combination of food abundance and temperature increase leading to size increases in mosquitoes. Across ectotherms, it has been said there is a need to reconcile the fact that when considering fitness, bigger is typically better, hotter is better (in the sense of having an optimal performance at higher temperatures), and hotter makes smaller^78^. By breaking TSR, fire ants can make bigger individuals in a shorter period of time as a function of increased temperature, potentially facilitating the production of an array of sizes including those requiring the most resources to make.

#### Nutritional Plasticity of Size and Developmental Timing

Looking at nutrition, a second E1 factor, we found that nutritional deficiency ranging from intermittent fasting to nutrient deprivation, influenced sizing and developmental timing to varying degrees (Fig. 3; Fig. 7, arrow 2). In the context of intermittent fasting, we found that individuals delayed metamorphosis and substantially extended their last instar, but they obtained the same final body size as optimally fed individuals (Fig. 3*A*, *B*, *C*). In the context of nutrient deprivation, where individuals obtained minimal nutrients, individuals extended their developmental duration and were substantially smaller than optimally fed individuals (Fig. S2). Our findings that nutritional variation can influence both developmental timing and sizing is consistent to what is known in other ant species. For example, in *Pheidole pallidula*, increasing levels of protein has been shown to increase growth and induce the production of large soldier and even supersoldiers^22,23^. In fire ants, colonies experiencing nutritional deprivation have a higher frequency of brood developing at lower temperatures^79^. In contrast, in well-fed colonies, there is a higher frequency of brood developing at higher temperatures^79^. It is possible that fire ants have evolved a superorganismal ability to integrate temperature and nutrition by cycling individual brood along their colony’s thermal gradient in a nutrition-dependent manner, impacting their demography.

Sociogenesis in ants is defined as the superorganismal development of the ant colony through its life-cycle as a society akin to the development and subsequent life-history of a solitary organism^19,80^. This life-cycle starts off with the queen experiencing nutritional deprivation^81^. The first workers laid by the queen, develop faster and smaller than any other workers for the rest of the life-cycle of the colony^33,82^. In fire ant colonies, as the colony develops and matures, it produces both a wide range of individual sizes including much larger individuals^19^. Whether small or big, quick or extended development, our findings suggest that the ability to respond to nutritional and thermal plasticity can together enable the tinkering of size and heterochrony, potentially enabling the stages of sociogenesis. Collectively, the elaboration of size range may be facilitated through the co-evolution of complex colony-level thermoregulation with the regulation of nutrient distribution in the colony.

### Mediators of Developmental Plasticity in fire ants

#### JH Signalling: Environmental Response and Influence on Sizing and Developmental Timing

Expanding to E2 (Endocrine mechanism), we found that JH influences the sizing of head and body, and developmental timing (Fig. 4*A-C*; Fig. 7, arrow 5). Based on extensive work done in fire ants, and ants of *Pheidole*, and *Camponotus*, our work reinforces the role of JH in regulating sizing and developmental timing^24,25,31,74,83,84^. Furthermore, we found that JH can change the allometry of developing *S. invicta* workers (Fig. 4*D*). Of note, the relative increase in the head-to-body slope, head size, and body size of JHA-treated individuals compared to controls, phenocopies the slope differences and morphospace exhibited by *Solenopsis geminata*, a species of fire ant that has a soldier caste^15,31^. We also found that temperature increase led to an increase in JH signalling, through *Kr-h1* activity (Fig. 2*E*; Fig. 7, arrow 3), yet nutritional variation (fed versus starved) did not influence JH signalling (Fig. 3*E*). Future work should characterize the role of JH outside of the early 4^th^ instar larval time period and further characterize, at shorter and longer timepoints, the influence of starvation on JH signalling.

#### Ecdysone Signalling: Environmental Response and Influence on Sizing and Developmental Timing

Looking at ecdysone, a second E2 factor, we found that ecdysone influences the sizing of head and body and developmental timing (Fig. 5*A, B, C*; Fig. 7, arrow 5). Surprisingly, ecdysone generates larger individuals, and these larger individuals reach their final size in a shorter time period, as compared to controls. This would indicate that ecdysone stimulates a faster growth rate to positively regulate size in fire ants as opposed to the extension of the duration of development that is regulated by JH. This is reminiscent of findings in honeybees where ecdysone promotes faster growth of larger (queens) individuals^85^. Furthermore, we found that increases in temperature led to an increase in head and body size, yet this was achieved through a shorter developmental duration (Fig. 2*-C*). This suggests that ecdysone controlling growth rate is potentially thermally plastic, which is further supported by our finding of nutrition positively regulating ecdysone signalling. Based on our JH and ecdysone experiments, and both pathways being positively influenced by temperature increase (Fig. 2*E, F*; Fig. 7, arrow 3), we therefore suggest that social insects may have independently co-opted JH and ecdysone signalling, and their synergy, to coordinate growth rates with developmental timing, facilitating the evolution of size variation within and between castes.

#### Histone (de)acetylation: Environmental Response and Influence on Sizing and Developmental Timing

Expanding to E3 (Epigenetic mechanism), we found for histone (de)acetylation that histone deacetylation modifiers (*Hdac1a* and *Hdac1b*) are associated with temperature and nutritional states that would enable larger sized individuals (Fig. 7, arrow 4). An increase in temperature as well as an increase in nutrition when comparing fed versus starvation both lead to increases in size and *Hdac1* activity (Fig. 2*A, B, H*; Fig. 3*H*; Fig. S2). In contrast, when inhibiting histone deacetylation, rather than obtaining smaller individuals, we obtained individuals with both larger head and body sizes (Fig. 6*A, B*; Fig. 7, arrow 6). Furthermore, HDACi treatments led to a shift in the distribution of individual sizes that span the upper size range, relative to DMSO controls (Fig. 6*A, B*). Changes in sizing through histone epigenetic mechanisms in fire ants is consistent with correlations found between caste and histone acetylation modifier activity in both the honeybee *A. mellifera* and the Florida Carpenter ant *C. floridanus*^26,86^. One discrepancy is that starvation (decreases size) led to a decrease in histone deacetylases, yet our functional inhibition of histone deacetylases increased size. Our *atg8* data suggests that autophagy was induced following HDACi and together with an abundance of food led to an increase in size. We hypothesize that for fire ants, the HDACi-induction of both ecdysone signalling and autophagy alongside an abundance of nutritional resources, leads to a positive regulation of size. Furthermore, our HDACi treatment generated larger individuals that obtained their increase in size independent of developmental timing, suggesting that their larger size was through an increase in the rate of growth rather than the duration of growth. Altogether, histone (de)acetylation may be an important integrator of plasticity and growth to coordinate developmental timing, growth rates, and size variation.

#### Histone (de)methylation: Environmental Response and Influence on Sizing and Developmental Timing

Looking at histone (de)methylation, a second E3 factor, we found that the histone demethylation modifier *Utx* is positively associated with nutrition-mediated increase in size (Fig. 3*K*; Fig. S2), and both *Utx* and the histone methylation modifier *Su(z)12* are negatively associated with temperature-mediated increase in size (Fig. 2*K, J*; Fig. 7, arrow 4). We found that by inhibiting histone methylation using an HMTi (GSK343), this causes a decrease in size (Fig. 6*F, G*; Fig. 7, arrow 6). Given that *Su(z)12* and *Utx* both target the Histone H3 lysine 27 residue, it could be suggested that histone demethylation (*Utx*) is analogous to HMTi activity (GSK343). Therefore, a decrease in size induced by HMTi (GSK343) treatment mirrors the lower temperature-induced increase in *Utx* expression and decrease in size (Fig. 3*K*). Lower temperatures leading to an upregulation of *Utx* in fire ants underlying size plasticity is reminiscent of lower temperature leading to an upregulation of its vertebrate ortholog *Kdm6b* in turtles, underlying the plasticity of sex^9^.

Our findings suggest that while histone acetylation positively regulates size, likely through the active transcription of developmental genes, surprisingly histone methylation (PRC2-repressive complex) also positively regulates size. While counterintuitive, there are instances in both a developmental and cancer context that perturbing the PRC2-repressive complex can lead to irreversible DNA methylation-mediated gene silencing. Specifically, chromatin bivalency: the simultaneous presence of both repressive and active histone methylation marks (H3K27me3 and H3K4me3, respectively), is a key poised state of chromatin, shielding the chromatic region from the repressive activity of DNA methylation^87–90^. Chromatin bivalency has recently been discovered to be functionally conserved in insects^91–95^. By perturbing the PRC2 complex, we may have disrupted this shielding state, enabling subsequent DNA methylation. It has been shown that increasing DNA methylation leads to a decrease in size of workers in the highly polymorphic ant *C. floridanus* as well as the monomorphic ants *M. pharaonis* and *L. humile*^26,27^. Furthermore, it has been shown in fire ants that the de novo DNA methyltransferase *Dnmt3* is downregulated in 4^th^ instar larvae^96^. Therefore, fire ants may need a shielding bivalent chromatin state to facilitate positive sizing plasticity akin to how this state is required for epigenetic plasticity more broadly, and by perturbing this state leads to a decrease in size similarly to the influence of DNA methylation in other species of ants.

#### Endocrine-Epigenetic Crosstalk Underlying Size Variation and Developmental Timing in Fire Ants

In the context of endocrine (E2)-epigenetic (E3) crosstalk, we found that juvenile hormone has a dramatic influence on phenotype (Fig. 4*A-D*). These phenotypic consequences are associated with our findings that JH signaling downregulates both histone methylation (*Su(z)12*) and demethylation (*Utx*) yet did not influence the activity of histone acetylation modifiers (Fig. 4*H, I*; Fig. 7, arrow 7). This is consistent with findings in flies and moths that *Su(z)12* is negatively regulated by JH^97^. Furthermore, we found that ecdysone had no effect on histone modifiers (Fig. 5*E-I*), and that there is a negative relationship between histone deacetylase activity and ecdysone signaling (Fig. 6*K*; Fig. 7, arrow 8) suggesting that ecdysone’s regulatory role in growth rate and sizing may be downstream of histone epigenetic mechanisms of size. Consistent with our results, histone acetylation has been shown to be crucial in regulating ecdysone response genes in a positive manner in a range of insects^98–101^. Throughout this study, an increase in ecdysone activity has been correlated with an increase in size in the contexts of thermal, nutritional, and histone modifier manipulations (Fig. 2, 3, 6) and has been demonstrated to increase size and shorten developmental timing (Fig. 5). Thus far, we know that HDAC activity has a negative influence on ecdysone activity in *Drosophila*^98^ and that both HDAC activity and ecdysone activity demonstrate respective negative and positive relationships with honeybee queen development^85,86,102,103^. Therefore, we suggest that histone (de)acetylation may regulate ecdysone activity to determine appropriate sizing across insects. Future work in social insects may better illuminate the complex dynamics between the regulation of both developmental timing and growth underlying sizing and size evolution across castes.

Collectively, endocrine and epigenetic factors, and the endocrine-epigenetic axis mediate thermal and nutritional plasticity of the extreme size variation within the worker caste system in fire ants. In complement to sociogenesis mentioned above, sociometry is the study of the various biological parameters of a colony at individual timepoints including allometry of individuals, reproduction, demography and ratio of castes, and colony size^19,104,105^. In our study, we measured how key sociometric values including size, allometry, and developmental timing of individuals is regulated by endocrine and epigenetic factors as well as the axis formed between the two. Tinkering with these molecular mechanisms and their interplay may shape sociometric values over time giving rise to the evolution and elaboration of the sociogenesis of a superorganism. While our study focused on developmental growth and timing, we propose that future work should investigate the endocrine-epigenetic axis across the sociometric landscape.

#### An E3 Approach Towards Understanding Developmental Plasticity and Phenotypic Variation

While temperature and nutrition have been extensively investigated in the context of phenotypic plasticity, by taking an E3 approach, we were still able to identify unexpected contributions of endocrinological and epigenetic mediators underlying thermal and nutritional plasticity. Beyond these two classic factors, there are a plethora of environmental variables that remain poorly investigated in the context of development and growth. By continuing to expand the scope of which abiotic and biotic cues are investigated, we hope that a more holistic picture will emerge regarding our understanding of how an organism develops in a dynamic environment. Finally, by extending this approach to a comparative framework, we can begin to understand how changes in environmental variation over time can lead to the evolution of novel phenotypes.

## MATERIALS & METHODS

### Ant husbandry

Monogynous *Solenopsis invicta* colonies originally collected in Florida, USA, were donated by the Abouheif lab. All colonies were maintained in incubators set to 25°C, 60% humidity and a 12:12 h light:dark cycle. Colonies were kept in plastic boxes with mesh-covered holes to allow for airflow, and fluon-coated walls to prevent insect climbing. The boxes were lined with test tubes filled with either water or sugar water and plugged with cotton to allow for enclosed living quarters for the ants. A layer of acetate paper covered the tubes to mimic a subterrestrial environment. Colonies were fed 3x/week with a diet of mealworms, fruit flies, and Bhatkar-Whitcomb diet^106^. All experiments except for ecdysone and nutritional treatments were performed on the same colony to allow for more consistent data production.

### Experimental replicate setup

From the *S. invicta* colonies, all experimental treatments were composed of worker ants and early 4^th^ instar larvae, isolated in a 4:1 ratio. E4 larvae were targeted because the last larval instar of *S. invicta* is the 4^th^ instar, which is a developmentally plastic period that allows for the highest growth potential to occur^26^. The early 4^th^ instar larvae have yet to experience these high levels of growth, enabling this developmental stage to respond to various experimental manipulations and observe their effects on growth and gene activity. Furthermore, the smallest observable workers were selected for each replicate. The smallest fire ant workers are known to be primarily dedicated to brood care^28^, thus ensuring sufficient care for developing larvae within replicates. Each replicate was separated into small plastic containers containing 1-2 small cotton-bound water tubes, mimicking the environment of their source colony. Larvae were measured using the Zeiss Axio Zoom. V16 microscope. Early 4^th^ instar larvae were categorized based on the outlined measurements of head width (0.26-0.33 mm) and body length (0.910-1.200 mm), and with the emergence partially sclerotized mouthparts, as described by Petralia & Vinson^30^. Furthermore, 3^rd^ instar larval head width ranged from 0.20-0.25 mm and body length from 0.60-0.90 mm, with the absence of mouthparts^30^. Late 4^th^ instar larval head width had the same range as the early 4^th^ instar individuals, but with a body length ranging from 1.50-1.82 mm^30^. Head width is the widest head capsule segment, while body length measures the top of the larval head to the base of its body (larvae were placed ventral side up and dorsal side down). These measurements were specific to the ‘minor worker’ destined larvae for *S. invicta*, meaning that these larval size ranges led to the smaller workers to be produced^30^. All of our experimental manipulations focused on the ‘minor worker’ destined larval size ranges. Furthermore, collected early 4^th^ instar larvae were separated into smaller (0.910-1.100 mm) and larger (1.101-1.200) groups and split evenly among each set of trials to allow for an equal distribution of ‘early’ and ‘late’ early 4^th^ instar larvae per treatment. With the exception of the nutritional experiments, all treatments were fed 3x/week with a diet 2 mealworms cut into 3-4 pieces and a cube of Bhatkar-Whitcomb diet, until final pupation.

### Morphological data collection

Treatments were checked every 2-4 days to record the progression of pupal distribution by dumping out glass tube contents and counting the pupae present, providing our developmental timing data. Counts could not be made daily without disturbing larval growth and promoting worker stress. Dumped pupae were then collected and the Zeiss Axio Zoom. V16 microscope was used for pupal measurements. The anterior of the pupae faces forward, with their dorsal segments facing upwards, as demonstrated by Rajakumar et al.^84^. From this, head width is measured using the widest head segment. Body length is measured from the top of the head to the base of the thorax, at a consistent physical landmark found among pupae, found in between the first and second petioles.

### RT-qPCR primer design

Our overall RT-qPCR primer design largely followed the MIQE guidelines^107–111^. Using the National Center for Biotechnology (NCBI) database, we designed 1-3 pairs of forward and reverse primers for each of our genes of interest^108^. The protein sequence of each gene was first found within the *D. melanogaster* database, given the high level of gene conservation in the robust fly genome. The protein sequence was then run through BLAST against the *S. invicta* database, in search for the corresponding sequence in fire ants^112^. Nucleotide sequences were then acquired and imported into the Geneious software, allowing for sequence annotations. Exon-exon junctions were located and placed along each sequence. Primers were then designed along the identified exon-exon junctions to avoid intron amplification^109^. Primer sizes ranged from 18-22 base pairs, with melting temperatures (T_m_) ranging from 55-65°C, with each pair being within 2 degrees of each other^108,110^. Their GC base pair content required a range of 50-60%^108^. Primers could not include four of the same bases in a row and we aimed for our total amplicon lengths to span 80-115 base pairs (bp), but could range from a minimum of 75 bp to a maximum of 150 bp^108–110^. Using Oligo Calc, self-complementarity and self-annealing of primers were verified and adjusted for. Each primer and amplicon were then NCBI blasted against *S. invicta*, to check for the specificity of the amplicon relative to the gene of interest^108,109^. A small E-value was required (∼0), which would indicate a decreased probability that the similarity between the sequences is due to chance^109^. Additionally, query coverage needed to be >90% for our genes of interest, while undesirable genes could not surpass 70%^109^. Finalized primer sequences were then ordered from ThermoFisher Scientific and tested using PCR/gel-electrophoresis from *S. invicta* cDNA, to select for the optimal primer pair for our targeted genes. Exceptionally, to ensure primers targeting paralogs, in this case *Hdac1a* and *Hdac1b*, were paralog-specific, their nucleotide sequences were aligned, and we designed primers in regions that were variable and lacked conservation between the two.

### RNA extraction and cDNA synthesis

RNA extraction and cDNA synthesis protocols were done as previously^74^. For each experimental manipulation with the intention for RT-qPCR analysis, 6 biological replicates were produced, each containing 5 larvae, for a total of 30 larvae per treatment group. Larvae were frozen within 1.5 mL centrifuge tubes at -80°C for a minimum of 24 hours prior to extraction and purification of RNA. This allowed larvae to be easily crushed and hand homogenized in 100 uL of Trizol. Tubes were filled up to 1 mL of Trizol and were left for 5 minutes to incubate at room temperature (RT). To separate the aqueous phase, 200 uL of chloroform was added with a 3-minute RT incubation, followed by a centrifugation at 12,000 x g at 4°C for 15 minutes. 500 uL of the aqueous phase was separated and combined with 500 uL of 99% ethanol. The Direct-zol RNA MicroPrep kit (Zymo Research) was then used to complete RNA extraction. To purify the RNA of any DNA contaminants, a Turbo DNase treatment (Invitrogen) was implemented. The purified RNA was then verified for its concentration and purify level using a nanodrop (ThermoFischer). From this, 500 ng (within 21 uL of solution) of cDNA was produced by first normalizing samples based on RNA concentrations, followed by using the Super Script IV kit (Invitrogen). To match the concentration of cDNA produced, RNA dilution of 500 ng in 21 uL of DEPC water was prepared, followed by a collective 20-fold dilution of both RNA and cDNA samples for RT-qPCR application.

### RT-qPCR

With the established primers for our genes of interest, we then used the extracted RNA and synthesized cDNA for RT-qPCR using relative quantification analysis. All RT-qPCR plates were run on a C1000 CFX96 RT-qPCR device, using SSO advanced SYBR green (BioRad). To understand how our genes of interest change in expression across treatments groups, we require a conserved gene that is highly developmentally and experimentally stable in its expression levels. The housekeeping gene *Rpl32* is a robust normalization gene, conserved in both solitary^113^ and social insects^114^, and previously used in ants^26^. With our housekeeping gene, relative fold changes were calculated using the ΔΔCT method^115^, to compare changes in gene expression between groups. First, CT values for the gene of interest and housekeeping gene were calculated, followed by the ΔCT value, which is the difference between the CT values of both genes. This normalized our sample, which was then subtracted from the ΔCT value of our control group, providing the ΔΔCT value. To provide biological relevance, 2^-ΔΔCT^ was calculated demonstrating relative fold changes of both treatment and control groups, for further statistical analysis.

### Temperature treatments

The three thermal conditions of low (25°C), medium (28.5°C), and high (32°C) were used to investigate effects of temperature variation on morphology. 25°C is the temperature at which colonies are normally reared, 32°C is the temperature for which *S. invicta* colonies are known to undergo maximum growth^33^, and 28.5°C provides an intermediate temperature. Three incubators were used to set each temperature, while humidity (60%) and photoperiod (12:12 h LD) were kept constant. Individual larvae were reared until pupation for morphological data collection. Three modified temperatures of a wider range including low (18°C), medium (25°C), and high (30.4°C) conditions, reflective of the range examined by Porter^33^, were used to rear larvae for 48 hours. This was followed by flash freezing at -80°C for RT-qPCR analysis. In both sets of temperatures, the 25°C groups were the controls, given that the source *S. invicta* colony is kept in a constant 25°C environment. All temperatures chosen were based on Porter’s study^33^, describing the ranges of temperatures that affect colony growth and developmental timing.

### Nutrition treatments

To investigate the effects of nutrition on developmental plasticity in *S. invicta*, replicates were prepared by first starving workers prior to early 4^th^ instar larvae being added to the experimental groups. Cassill & Tschinkel^116^ implemented a 48-hour worker starvation period, presumably to deplete workers of any residual nutrients. However, our trials suffered mass deaths from this method. Therefore, we starved workers for 24 hours prior to being added to replicate boxes for both groups. Both conditions were fed the same diet, consisting of 2 mealworms and a cube of Bhatkar-Whitcomb diet^106^, but differed in the frequency of feeding, where the controls were fed daily, while larvae experiencing caloric deficit were intermittently fasted, being fed every 4 days. Individual larvae were reared until pupation for morphological data collection. Furthermore, to determine the effects of nutrient deprivation on underlying endocrine and epigenetic activities, another set of replicates were prepared, where the control group was fed daily, while the treatment group was starved of any nutrients for 4 days (96 hours). Based on our findings from the nutrient deprivation setup, where there is a high mortality of adult workers caring for the experimental larvae, our 4-day starvation setup for quantifying the influence of nutritional variation on endocrine and epigenetic modifier expression levels, adult workers were not pre-starved to ensure adult workers could survive during the 96hr experimental period to care for the larvae being investigated. Larvae were then collected and flash frozen at -80°C for RNA extraction and RT-qPCR analysis. Finally, another set of treatments were arranged to investigate the effects of a high sugar diet, while being protein (among other nutrients) deficient. Controls were supplied with a saturated solution of sucrose and fed regularly every 2 days with two mealworms and a Bhatkar-Whitcomb diet, while the treatment only received the sucrose solution. Individual larvae were reared until pupation for morphological data collection.

### Endocrine manipulations

Hormonal treatments to target the JH pathway implemented the JH analogue methoprene (CAT: 33375; Millipore Sigma). 5 mg/mL aliquots were made using acetone as the solvent and were further diluted to 2.5 mg/mL. Treatments were carried out on early 4^th^ instar larvae, as previously described. Larvae were placed on their dorsal side on a wet Kimtech tissue. 1 ul of acetone (control) or methoprene (treatment) was applied topically on individual larval abdomens, with the high volatility of acetone allowing for rapid cuticular absorption^31^. Individual larvae were reared until pupation for morphological data collection or were collected after 24 hours post treatment and flash frozen at -80°C for RNA extraction and RT-qPCR analysis.

For ecdysone treatments, 20-hydroxyecdysone (20E) (CAT: H5142; Millipore Sigma) was dissolved in a 99% ethanol solvent, and 1 mg/mL aliquots were used for microinjection experimental manipulations, using techniques described by Bear et al.^117^. Both the 99% ethanol control and 20E treatment were dissolved in a 1x phosphate-buffered saline (PBS) solution, at a 1:9 ratio, to allow for optimal hormonal delivery. Individual early 4^th^ instar larvae were microinjected with either 99% ethanol or with 1 mg/mL 20E, using a combination of a Zeiss SteREO Discovery V8 microscope with an Eppendorf CellTram 4r Oil microinjector and a Narishige micromanipulator. A Stutter Instrument P-97 needle puller was used to prepare needles for larval injection. Larval bodies were placed ventral-up, and their cuticles were pierced at the lateral midline, which is a fatty body section, and in between larval segments, to minimize fatal wounds. Individual larvae were reared until pupation for morphological data collection or were collected after 24 hours post treatment and flash frozen at -80°C for RNA extraction and RT-qPCR analysis.

### Pharmacological histone modification inhibitors

The pan-histone deacetylase inhibitor trichostatin-A (TSA) (CAT: T1952; Millipore Sigma) was acquired at 5 mM, dissolved in a dimethyl sulfoxide (DMSO) solvent for experimental manipulation as done previously in ants^46^. Individual early 4^th^ instar larvae were microinjected using the same methodology as in the 20E treatments. However, given the high mortality rate of DMSO dissolved treatments, the proportion of the PBS vehicle present was increased to mitigate the lethality of DMSO. Therefore, DMSO controls and TSA treatments were dissolved in 1x PBS at a 0.5:9.5 ratio. The histone methyltransferase inhibitor GSK343 (CAT: SML0766; Millipore Sigma), which targets a component of the PRC2 silencing complex, was dissolved in DMSO at a concentration of 5 mM as done previously in insects and nematodes^64,118^. Methodology for PBS dilution and microinjection followed the same protocol as the TSA treatments. Individual larvae were reared until pupation for morphological data collection or were collected after 48 hours post treatment ^46^ and flash frozen at -80°C for RNA extraction and RT-qPCR analysis.

### Statistical analyses

All datasets were tested for normality using the Shapiro-Wilk test, while the developmental timing results were tested using a combination of the Shapiro-Wilk and D’Agostino & Pearson tests, given the several identical values found within the developmental timing datasets. If following normal distribution, the means of head width, body length, developmental timing, and RT-qPCR relative fold changes were respectively compared using one-way ANOVA analyses (followed by Tukey’s multiple comparison test) for treatments that were experimenting with three parameters (natural expression and temperature treatments), while unpaired two-tailed Student’s *t*-tests were used for treatments with two parameters (nutrition, endocrine, and epigenetic). If the dataset is not normally distributed, a Kruskal-Wallis test (followed by Dunn’s multiple comparison test) was used rather than an ANOVA, while a Mann-Whitney test was used instead of a Student’s *t*-tests. A one-tailed *t*-test was used to assess the effect of JHA on *Su(z)12* (Fig. 4*I*), because it has been established that *Su(z)12* is negatively regulated by JH signalling and specifically by JHA treatments in *Bombyx* and *Drosophila*^97^. Furthermore, a Welch’s *t*-test was used in the analysis of TSA treatments on head and body sizes (Fig. 6*A, B*), given their significance in F test variance. Scatterplots were log transformed and both slopes and (when possible) y-intercepts were compared between groups. All analyses were done using GraphPad Prism.

## ACKNOLWEDGEMENTS

We thank Ehab Abouheif for *Solenopsis invicta* colonies; A. Rajakumar, H. MacMillan, G. Stefanelli, and Rajakumar Lab members for comments. RR acknowledges funding support of the Natural Sciences and Engineering Research Council of Canada (NSERC) [RGPAS-2021-00006; RGPIN-2021-04399; RTI-2021-00710]; Canadian Foundation for Innovation and the Ontario Research Fund [40142].

## AUTHOR CONTRIBUTIONS

Conceptualization and experimental design: N.B., R.R.

Experimentation: N.B., A.D., V.T., K.F., R.R.

Visualization and analysis: N.B., V.T.

Funding acquisition: R.R.

Supervision: R.R.

Writing – original draft: N.B., R.R.

Writing – review & editing: N.B., R.R.

## SUPPLEMENTAL INFORMATION

**Fig. S1.**
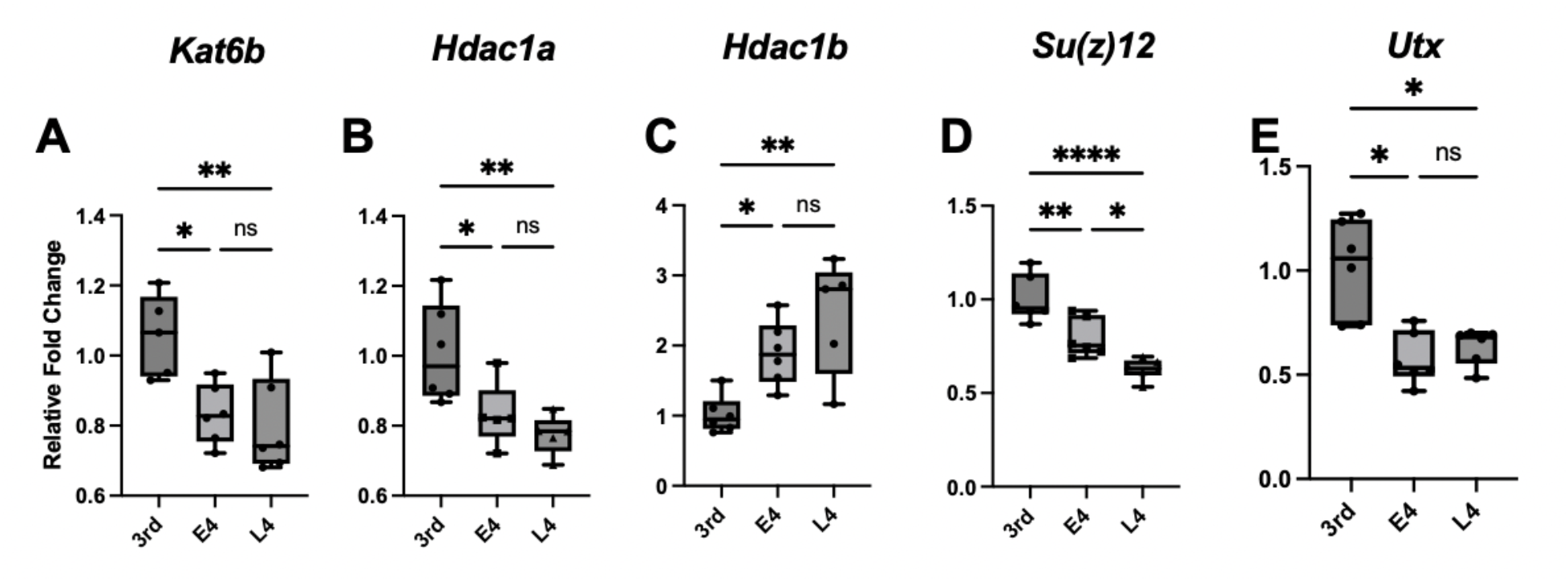
The expression levels of targeted histone modifier toolkit genes across key larval stages of the high frequency small worker. qPCR analyses were used to compare the relative expression levels of target genes in developing 3^rd^, early 4^th^ (E4) and late 4^th^ (L4) instar larvae. One-way ANOVA analyses were used to compare expression differences in **(A)** *Kat6b*, **(B)** *Hdac1a*, **(C)** *Hdac1b*, **(D)** *Su(z)12*, and **(E)** *Utx*. Bars represent means, with standard deviation (SD) indicated with error bars. Values for statistical significance of ANOVA and Kruskal-Wallis tests include \**p*<0.05, \*\**p*<0.01, \*\*\**p*<0.001, and \*\*\*\**p*<0.0001. Sample size for each group ranges from 5-6 biological replicates, each containing 5 individual larvae.

**Fig. S2.**
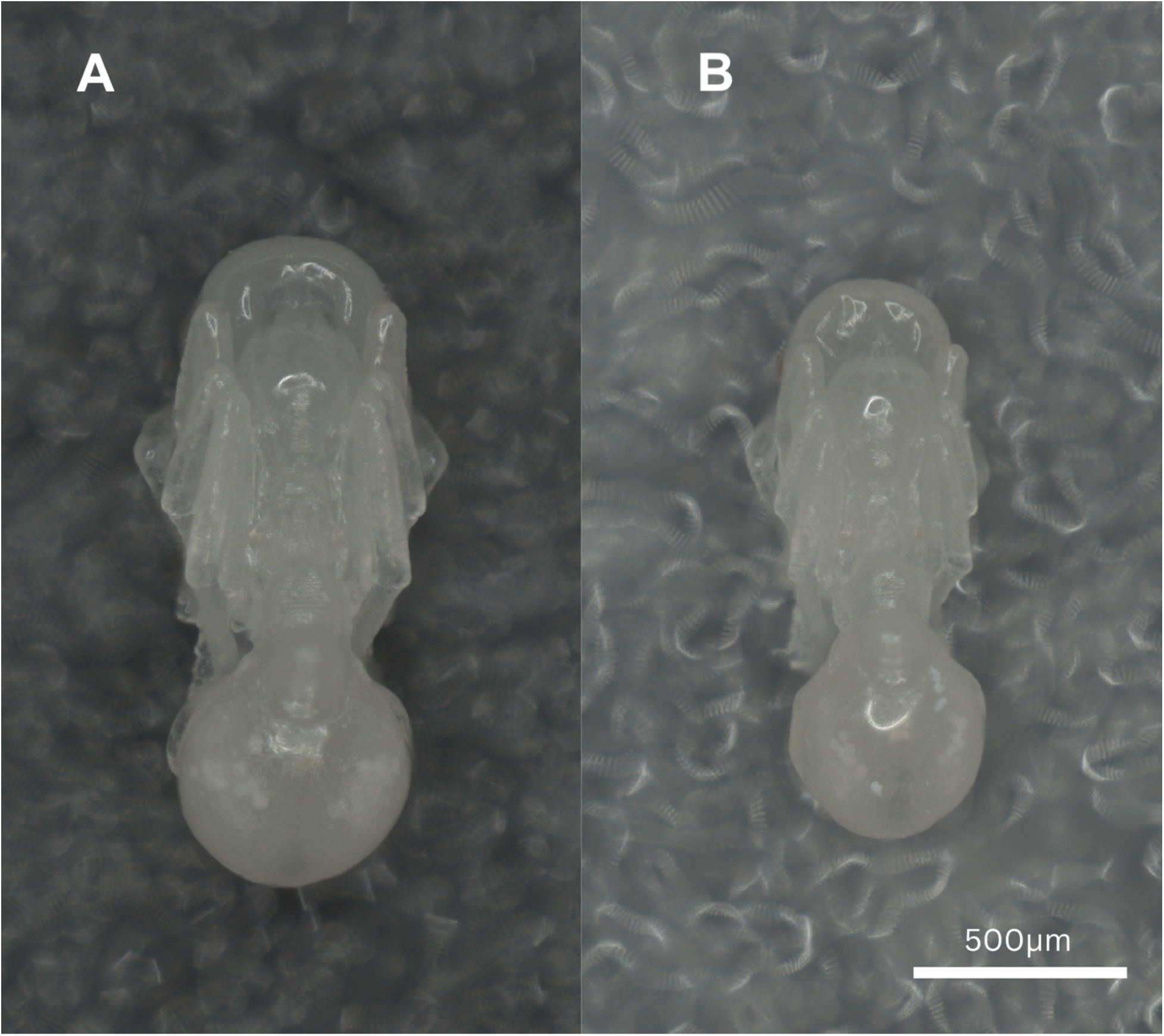
Comparison between the average sized fed and sugar-diet individual *S. invicta* pupae. **(A)** The average sized individual from 47 acquired pupae in a fed experiment. **(B)** The average sized individual from 4 acquired pupae in the sugar-diet (starved) experiment.

**Table S1:** Summary chart outlining the effects of each experimental manipulation on sizing, developmental timing, and allometry. Arrows indicate direction of change and number of arrows are representative of the significance level, based on statistical analyses: ↑/ ↓ represents \**p*<0.05; ↑↑/ ↓↓ represents \*\**p*<0.01; ↑↑↑/ ↓↓↓ represents \*\*\**p*<0.001; ↑↑↑↑/ ↓↓↓↓ represents \*\*\*\**p*<0.0001. Equal symbols demonstrate stability in gene expression.

|  | Head Width | Body Length | Dev Timing | XY changes |
| --- | --- | --- | --- | --- |
| Increasing temperatures | ↑ at 28.5°C | ↑ at 28.5°C | ↓↓↓↓ at 28.5°C and 32°C | ↓↓ in y-intercept at 28.5°C |
| Intermittent fasting | No change | No change | ↑↑↑↑ | No change |
| Methoprene | ↑↑↑↑ | ↑↑↑↑ | ↑↑↑↑ | ↑↑ in slope |
| 20E | ↑↑↑ | ↑↑↑↑ | ↓ | No change |
| TSA | ↑↑ | ↑ | No change | No change |
| GSK343 | ↓↓↓↓ | ↓↓↓↓ | No change | No change |

**Table S2:**
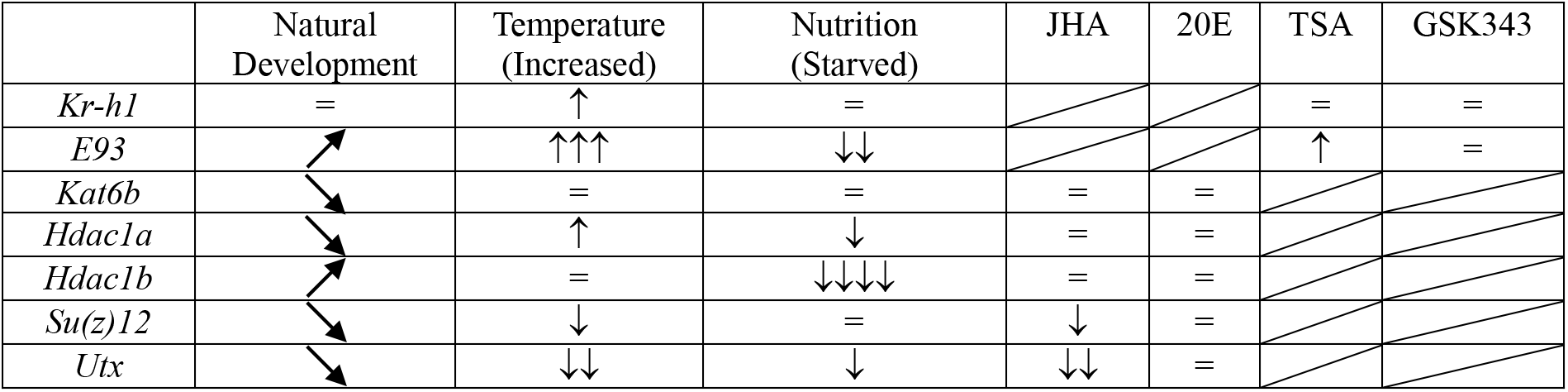
Summary chart outlining the responsiveness of JH/ecdysone signalling and of histone modifiers after various experimental manipulations. Diagonal arrows for Natural Development indicate the direction of change, leading to an overall increase or decrease in gene expression. Arrows indicate direction of change and number of arrows are representative of the significance level, based on statistical analyses: ↑/ ↓ represents \**p*<0.05; ↑↑/ ↓↓ represents \*\**p*<0.01; ↑↑↑/ ↓↓↓ represents \*\*\**p*<0.001; ↑↑↑↑/ ↓↓↓↓ represents \*\*\*\**p*<0.0001. Equal symbols demonstrate stability in gene expression.

**Table S3:**
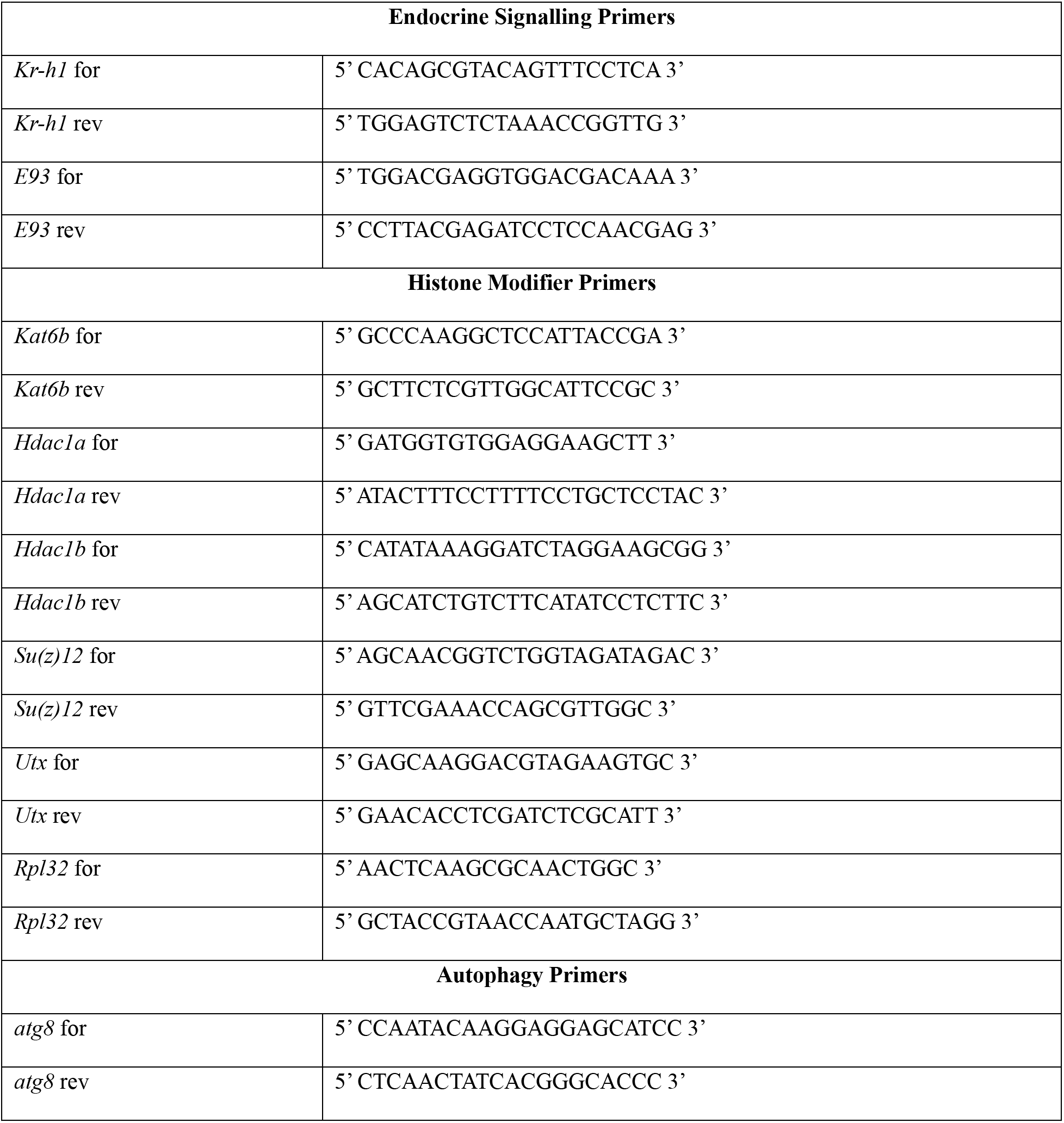
List of both forward (for) and reverse (rev) primers designed for RT-qPCR genes.

## REFERENCES

1. Gilbert, S.F., and Pfennig, D.W. (2026). Eco-evo-devo: The Environmental Regulation of Development, Health, and Evolution (Oxford University Press).

2. West-Eberhard, M.J. (1989). Phenotypic plasticity and the origins of diversity. Annu. Rev. Ecol. Syst. 20, 249–278.

3. West-Eberhard, M.J. (2003). Developmental plasticity and evolution (Oxford University Press).

4. Pfennig, D.W. (2021). Phenotypic plasticity & evolution: causes, consequences, controversies (Taylor & Francis).

5. Seebacher, F., and Little, A.G. (2026). Mechanisms underlying phenotypic plasticity in response to environmental change. Biol. Rev. 101, 713–734.

6. Miner, B.G., Sultan, S.E., Morgan, S.G., Padilla, D.K., and Relyea, R.A. (2005). Ecological consequences of phenotypic plasticity. Trends Ecol. Evol. 20, 685–692.

7. Crews, D., Bergeron, J.M., Bull, J.J., Flores, D., Tousignant, A., Skipper, J.K., and Wibbels, T. (1994). Temperature-dependent sex determination in reptiles: Proximate mechanisms, ultimate outcomes, and practical applications. Dev. Genet. 15, 297–312.

8. Crews, D., and Bergeron, J. (1994). Role of reductase and aromatase in sex determination in the red-eared slider (Trachemys scripta), a turtle with temperature-dependent sex determination. J. Endocrinol. 143, 279–289.

9. Ge, C., Ye, J., Weber, C., Sun, W., Zhang, H., Zhou, Y., Cai, C., Qian, G., and Capel, B. (2018). The histone demethylase KDM6B regulates temperature-dependent sex determination in a turtle species. Science 360, 645–648.

10. Wirtz, P., and Beetsma, J. (1972). Induction of caste differentiation in the honeybee (Apis mellifera) by juvenile hormone. Entomol. Exp. Appl. 15, 517–520.

11. Goewie, E., and Beetsma, J. (1976). Induction of caste differentiation in the honey bee (Apis mellifera L.) after topical application of JH-III. Proc. K. Ned. Akad. Van Wet. C 79, 466–469.

12. Kucharski, R., Maleszka, J., Foret, S., and Maleszka, R. (2008). Nutritional control of reproductive status in honeybees via DNA methylation. Science 319, 1827–1830.

13. Zhu, K., Liu, M., Fu, Z., Zhou, Z., Kong, Y., Liang, H., Lin, Z., Luo, J., Zheng, H., and Wan, P. (2017). Plant microRNAs in larval food regulate honeybee caste development. PLoS Genet. 13, e1006946.

14. Lewontin, R.C. (2000). The triple helix: Gene, organism, and environment (Harvard University Press).

15. Tschinkel, W.R. (2006). The Fire Ants (Harvard University Press).

16. Cuthbert, R.N., Diagne, C., Haubrock, P.J., Turbelin, A.J., and Courchamp, F. (2022). Are the “100 of the world’s worst” invasive species also the costliest? Biol. Invasions 24, 1895–1904.

17. Mirenda, J.T., and Vinson, S.B. (1981). Division of labour and specification of castes in the red imported fire ant Solenopsis invicta Buren. Anim. Behav. 29, 410–420.

18. Hölldobler, B., and Wilson, E.O. (2009). The superorganism: the beauty elegance and strangeness of insect societies (WW Norton & Company).

19. Tschinkel, W.R. (2011). Back to basics: sociometry and sociogenesis of ant societies (Hymenoptera: Formicidae). Myrmecol. News 14, 49–54.

20. Hölldobler, B., and Wilson, E.O. (1990). The ants (Belknap Press of Harvard University Press).

21. Abouheif, E., and Wray, G.A. (2002). Evolution of the gene network underlying wing polyphenism in ants. Science 297, 249–252.

22. Goetsch, W. (1937). Die Entstehung der „Soldaten “im Ameisenstaat. Naturwissenschaften 25, 803–808.

23. Passera, L. (1974). Différenciation des soldats chez la Fourmi Pheidole pallidula Nyl. (Formicidae Myrmicinae). Insectes Sociaux 21, 71–86.

24. Wheeler, D.E., and Nijhout, H.F. (1981). Soldier determination in ants: new role for juvenile hormone. Science 213, 361–363.

25. Rajakumar, R., San Mauro, D., Dijkstra, M.B., Huang, M.H., Wheeler, D.E., Hiou-Tim, F., Khila, A., Cournoyea, M., and Abouheif, E. (2012). Ancestral developmental potential facilitates parallel evolution in ants. Science 335, 79–82.

26. Alvarado, S., Rajakumar, R., Abouheif, E., and Szyf, M. (2015). Epigenetic variation in the Egfr gene generates quantitative variation in a complex trait in ants. Nat. Commun. 6, 6513.

27. Renard, T., Gueydan, C., and Aron, S. (2022). DNA methylation and expression of the egfr gene are associated with worker size in monomorphic ants. Sci. Rep. 12, 21228.

28. Wilson, E.O. (1978). Division of labor in fire ants based on physical castes (Hymenoptera: Formicidae: Solenopsis). J. Kans. Entomol. Soc., 615–636.

29. Cassill, D.L., and Tschinkel, W.R. (1999). Task selection by workers of the fire ant Solenopsis invicta. Behav. Ecol. Sociobiol. 45, 301–310.

30. Petralia, R.S., and Vinson, S.B. (1979). Developmental Morphology of Larvae and Eggs of the Imported Fire Ant, Solenopsis invicta. Ann. Entomol. Soc. Am. 72, 472–484.

31. Wheeler, D.E. (1990). The developmental basis of worker polymorphism in fire ants. J. Insect Physiol. 36, 315–322.

32. Hanna, L., Lamouret, T., Poças, G.M., Mirth, C.K., Moczek, A.P., Nijhout, F.H., and Abouheif, E. (2023). Evaluating old truths: Final adult size in holometabolous insects is set by the end of larval development. J. Exp. Zoolog. B Mol. Dev. Evol. 340, 270–276.

33. Porter, S.D. (1988). Impact of temperature on colony growth and developmental rates of the ant, Solenopsis invicta. J. Insect Physiol. 34, 1127–1133.

34. Atkinson, D. (1994). Temperature and organism size-a biological law for ectotherms? Adv Ecol Res 25, 1–58.

35. Truman, J.W., and Riddiford, L.M. (1999). The origins of insect metamorphosis. Nature 401, 447–452.

36. Edgar, B.A. (2006). How flies get their size: genetics meets physiology. Nat. Rev. Genet. 7, 907–916.

37. Belles, X. (2020). Krüppel homolog 1 and E93: The doorkeeper and the key to insect metamorphosis. Arch. Insect Biochem. Physiol. 103, e21609.

38. Nicholson, T.B., Veland, N., and Chen, T. (2015). Writers, Readers, and Erasers of Epigenetic Marks. In Epigenetic Cancer Therapy (Elsevier), pp. 31–66.

39. Marmorstein, R., and Zhou, M.-M. (2014). Writers and readers of histone acetylation: structure, mechanism, and inhibition. Cold Spring Harb. Perspect. Biol. 6, a018762.

40. Seto, E., and Yoshida, M. (2014). Erasers of histone acetylation: the histone deacetylase enzymes. Cold Spring Harb. Perspect. Biol. 6, a018713.

41. Ringrose, L., and Paro, R. (2004). Epigenetic regulation of cellular memory by the Polycomb and Trithorax group proteins. Annu Rev Genet 38, 413–443.

42. Romiguier, J., Borowiec, M.L., Weyna, A., Helleu, Q., Loire, E., La Mendola, C., Rabeling, C., Fisher, B.L., Ward, P.S., and Keller, L. (2022). Ant phylogenomics reveals a natural selection hotspot preceding the origin of complex eusociality. Curr. Biol. 32, 2942–2947.e4.

43. Vizueta, J., Xiong, Z., Ding, G., Larsen, R.S., Ran, H., Gao, Q., Stiller, J., Dai, W., Jiang, W., and Zhao, J. (2025). Adaptive radiation and social evolution of the ants. Cell.

44. Gospocic, J., Glastad, K.M., Sheng, L., Shields, E.J., Berger, S.L., and Bonasio, R. (2021). Kr-h1 maintains distinct caste-specific neurotranscriptomes in response to socially regulated hormones. Cell 184, 5807–5823.

45. Perez, R., De Souza Araujo, N., Defrance, M., and Aron, S. (2021). Molecular adaptations to heat stress in the thermophilic ant genus *Cataglyphis*. Mol. Ecol. 30, 5503–5516.

46. Simola, D.F., Graham, R.J., Brady, C.M., Enzmann, B.L., Desplan, C., Ray, A., Zwiebel, L.J., Bonasio, R., Reinberg, D., Liebig, J., et al. (2016). Epigenetic (re)programming of caste-specific behavior in the ant *Camponotus floridanus*. Science 351, aac6633.

47. Brian, M.V. (1956). Studies of caste differentiation in Myrmica rubra L. 4. Controlled larval nutrition. Insectes Sociaux 3, 369–394.

48. Dussutour, A., and Simpson, S.J. (2009). Communal Nutrition in Ants. Curr. Biol. 19, 740–744.

49. Csata, E., and Dussutour, A. (2019). Nutrient regulation in ants (Hymenoptera: Formicidae): a review.

50. Mirth, C.K., and Riddiford, L.M. (2007). Size assessment and growth control: how adult size is determined in insects. BioEssays 29, 344–355.

51. Frederik Nijhout, H. (2013). Arthropod developmental endocrinology. In Arthropod Biology and Evolution: Molecules, Development, Morphology (Springer), pp. 123–148.

52. Brian, M.V. (1974). Caste differentiation in Myrmica rubra: The rôle of hormones. J. Insect Physiol. 20, 1351–1365.

53. Suzzoni, J., Passera, L., and Strambi, A. (1980). Ecdysteroid titre and caste determination in the ant, Pheidole pallidula (Nyl.)(Hymenoptera: Formicidae). Experientia 36, 1228–1229.

54. Schwander, T., Humbert, J.-Y., Brent, C.S., Cahan, S.H., Chapuis, L., Renai, E., and Keller, L. (2008). Maternal effect on female caste determination in a social insect. Curr. Biol. 18, 265–269.

55. Cahan, S.H., Graves, C.J., and Brent, C.S. (2011). Intergenerational effect of juvenile hormone on offspring in Pogonomyrmex harvester ants. J. Comp. Physiol. B 181, 991–999.

56. Colombani, J., Bianchini, L., Layalle, S., Pondeville, E., Dauphin-Villemant, C., Antoniewski, C., Carré, C., Noselli, S., and Léopold, P. (2005). Antagonistic actions of ecdysone and insulins determine final size in Drosophila. Science 310, 667–670.

57. McBrayer, Z., Ono, H., Shimell, M., Parvy, J.-P., Beckstead, R.B., Warren, J.T., Thummel, C.S., Dauphin-Villemant, C., Gilbert, L.I., and O’Connor, M.B. (2007). Prothoracicotropic hormone regulates developmental timing and body size in Drosophila. Dev. Cell 13, 857–871.

58. Tatun, N., Kumdi, P., Tungjitwitayakul, J., and Sakurai, S. (2018). Effects of 20-hydroxyecdysone on the development and morphology of the red fl our beetle, Tribolium castaneum (Coleoptera: Tenebrionidae). Eur. J. Entomol. 115, 424–431.

59. Jenuwein, T., and Allis, C.D. (2001). Translating the histone code. Science 293, 1074–1080.

60. Yoshida, M., Kijima, M., Akita, M., and Beppu, T. (1990). Potent and specific inhibition of mammalian histone deacetylase both in vivo and in vitro by trichostatin A. J. Biol. Chem. 265, 17174–17179.

61. George, S., Gaddelapati, S.C., and Palli, S.R. (2019). Histone deacetylase 1 suppresses Krüppel homolog 1 gene expression and influences juvenile hormone action in Tribolium castaneum. Proc. Natl. Acad. Sci. 116, 17759–17764.

62. Glastad, K.M., Graham, R.J., Ju, L., Roessler, J., Brady, C.M., and Berger, S.L. (2020). Epigenetic regulator CoREST controls social behavior in ants. Mol. Cell 77, 338–351.

63. Choppin, M., Feldmeyer, B., and Foitzik, S. (2021). Histone acetylation regulates the expression of genes involved in worker reproduction in the ant Temnothorax rugatulus. BMC Genomics 22, 871.

64. Werner, M.S., Loschko, T., King, T., Reich, S., Theska, T., Franz-Wachtel, M., Macek, B., and Sommer, R.J. (2023). Histone 4 lysine 5/12 acetylation enables developmental plasticity of Pristionchus mouth form. Nat. Commun. 14, 2095.

65. Berger, S.L. (2007). The complex language of chromatin regulation during transcription. Nature 447, 407–412.

66. Chang, Y.-Y., and Neufeld, T.P. (2010). Autophagy takes flight in Drosophila. FEBS Lett. 584, 1342–1349.

67. Glick, D., Barth, S., and Macleod, K.F. (2010). Autophagy: cellular and molecular mechanisms. J. Pathol. 221, 3–12.

68. Zhang, J., Ng, S., Wang, J., Zhou, J., Tan, S.-H., Yang, N., Lin, Q., Xia, D., and Shen, H.-M. (2015). Histone deacetylase inhibitors induce autophagy through FOXO1-dependent pathways. Autophagy 11, 629–642.

69. Kirisako, T., Baba, M., Ishihara, N., Miyazawa, K., Ohsumi, M., Yoshimori, T., Noda, T., and Ohsumi, Y. (1999). Formation process of autophagosome is traced with Apg8/Aut7p in yeast. J. Cell Biol. 147, 435–446.

70. Wang, P., Nolan, T.M., Yin, Y., and Bassham, D.C. (2020). Identification of transcription factors that regulate ATG8 expression and autophagy in Arabidopsis. Autophagy 16, 123–139.

71. Galves, M., Sperber, M., Amer-Sarsour, F., Elkon, R., and Ashkenazi, A. (2023). Transcriptional profiling of the response to starvation and fattening reveals differential regulation of autophagy genes in mammals. Proc. R. Soc. B Biol. Sci. 290, 20230407.

72. Margueron, R., and Reinberg, D. (2011). The Polycomb complex PRC2 and its mark in life. Nature 469, 343–349.

73. Ding, M., Zhang, H., Li, Z., Wang, C., Chen, J., Shi, L., Xu, D., and Gao, Y. (2015). The polycomb group protein enhancer of zeste 2 is a novel therapeutic target for cervical cancer. Clin. Exp. Pharmacol. Physiol. 42, 458–464.

74. MacMillan, O., Singer, J., Perrakis, S., Craig, A., Ntanga, D., Qiu, D., and Rajakumar, R. (2025). Juvenile Hormone Signalling Underlies the Switchpoint and Differentiation of Soldiers in *Camponotus floridanus*. Preprint.

75. La Richelière, F., Muñoz, G., Guénard, B., Dunn, R.R., Economo, E.P., Powell, S., Sanders, N.J., Weiser, M.D., Abouheif, E., and Lessard, J.-P. (2022). Warm and arid regions of the world are hotspots of superorganism complexity. Proc. R. Soc. B Biol. Sci. 289, 20211899.

76. Reiskind, M.H., and Zarrabi, A.A. (2012). Is bigger really bigger? Differential responses to temperature in measures of body size of the mosquito, Aedes albopictus. J. Insect Physiol. 58, 911–917.

77. Penick, C.A., and Tschinkel, W. (2008). Thermoregulatory brood transport in the fire ant, Solenopsis invicta. Insectes Sociaux 55, 176–182.

78. Kingsolver, J., and Huey, R. (2008). Size, temperature, and fitness: three rules. Evol. Ecol. Res. 10, 251–268.

79. Porter, S.D., and Tschinkel, W.R. (1993). Fire ant thermal preferences: behavioral control of growth and metabolism. Behav. Ecol. Sociobiol. 32, 321–329.

80. Wilson, E.O. (1985). The sociogenesis of insect colonies. Science 228, 1489–1495.

81. Matte, A., and Billen, J. (2021). Flight muscle histolysis in Lasius niger queens. Asian Myrmecol. 13.

82. Porter, S.D., and Tschinkel, W.R. (1986). Adaptive value of nanitic workers in newly founded red imported fire ant colonies (Hymenoptera: Formicidae). Ann. Entomol. Soc. Am. 79, 723–726.

83. Wheeler, D.E., and Nijhout, H.F. (1983). Soldier determination in Pheidole bicarinata: effect of methoprene on caste and size within castes. J. Insect Physiol. 29, 847–854.

84. Rajakumar, R., Koch, S., Couture, M., Favé, M.-J., Lillico-Ouachour, A., Chen, T., De Blasis, G., Rajakumar, A., Ouellette, D., and Abouheif, E. (2018). Social regulation of a rudimentary organ generates complex worker-caste systems in ants. Nature 562, 574–577.

85. Kamakura, M. (2011). Royalactin induces queen differentiation in honeybees. Nature 473, 478–483.

86. Spannhoff, A., Kim, Y.K., Raynal, N.J. -M, Gharibyan, V., Su, M., Zhou, Y., Li, J., Castellano, S., Sbardella, G., Issa, J.J., et al. (2011). Histone deacetylase inhibitor activity in royal jelly might facilitate caste switching in bees. EMBO Rep. 12, 238–243.

87. Eckersley-Maslin, M.A., Parry, A., Blotenburg, M., Krueger, C., Ito, Y., Franklin, V.N.R., Narita, M., D’Santos, C.S., and Reik, W. (2020). Epigenetic priming by Dppa2 and 4 in pluripotency facilitates multi-lineage commitment. Nat. Struct. Mol. Biol. 27, 696–705.

88. Gretarsson, K.H., and Hackett, J.A. (2020). Dppa2 and Dppa4 counteract de novo methylation to establish a permissive epigenome for development. Nat. Struct. Mol. Biol. 27, 706–716.

89. Kumar, D., Cinghu, S., Oldfield, A.J., Yang, P., and Jothi, R. (2021). Decoding the function of bivalent chromatin in development and cancer. Genome Res. 31, 2170–2184.

90. Weigert, R., Hetzel, S., Bailly, N., Haggerty, C., Ilik, I.A., Yung, P.Y.K., Navarro, C., Bolondi, A., Kumar, A.S., and Anania, C. (2023). Dynamic antagonism between key repressive pathways maintains the placental epigenome. Nat. Cell Biol. 25, 579–591.

91. Schertel, C., Albarca, M., Rockel-Bauer, C., Kelley, N.W., Bischof, J., Hens, K., Van Nimwegen, E., Basler, K., and Deplancke, B. (2015). A large-scale, in vivo transcription factor screen defines bivalent chromatin as a key property of regulatory factors mediating Drosophila wing development. Genome Res. 25, 514–523.

92. Kang, H., Jung, Y.L., McElroy, K.A., Zee, B.M., Wallace, H.A., Woolnough, J.L., Park, P.J., and Kuroda, M.I. (2017). Bivalent complexes of PRC1 with orthologs of BRD4 and MOZ/MORF target developmental genes in Drosophila. Genes Dev. 31, 1988–2002.

93. Akmammedov, A., Geigges, M., and Paro, R. (2019). Bivalency in Drosophila embryos is associated with strong inducibility of Polycomb target genes. Fly (Austin) 13, 42–50.

94. Cheng, Q., and Xie, H. (2022). Genome-wide analysis of bivalent histone modifications during Drosophila embryogenesis. genesis 60, e23502.

95. Bamgbose, G., and Tulin, A. (2024). PARP-1 is a transcriptional rheostat of metabolic and bivalent genes during development. Life Sci. Alliance 7, e202302369.

96. Kay, S., Skowronski, D., and Hunt, B.G. (2018). Developmental DNA methyltransferase expression in the fire ant Solenopsis invicta. Insect Sci. 25, 57–65.

97. Yang, Y., Zhao, T., Li, Z., Qian, W., Peng, J., Wei, L., Yuan, D., Li, Y., Xia, Q., and Cheng, D. (2021). Histone H3K27 methylation–mediated repression of Hairy regulates insect developmental transition by modulating ecdysone biosynthesis. Proc. Natl. Acad. Sci. 118, e2101442118.

98. Kirilly, D., Wong, J.J.L., Lim, E.K.H., Wang, Y., Zhang, H., Wang, C., Liao, Q., Wang, H., Liou, Y.-C., and Wang, H. (2011). Intrinsic epigenetic factors cooperate with the steroid hormone ecdysone to govern dendrite pruning in Drosophila. Neuron 72, 86–100.

99. Bodai, L., Zsindely, N., Gáspár, R., Kristó, I., Komonyi, O., and Boros, I.M. (2012). Ecdysone Induced Gene Expression Is Associated with Acetylation of Histone H3 Lysine 23 in Drosophila melanogaster. PLoS ONE 7, e40565.

100. Roy, A., and Palli, S.R. (2018). Epigenetic modifications acetylation and deacetylation play important roles in juvenile hormone action. BMC Genomics 19, 934.

101. Cheng, D., Dong, Z., Lin, P., Shen, G., and Xia, Q. (2022). Transcriptional Activation of Ecdysone-Responsive Genes Requires H3K27 Acetylation at Enhancers. Int. J. Mol. Sci. 23, 10791.

102. Polsinelli, G.A., and Yu, H.D. (2018). Regulation of histone deacetylase 3 by metal cations and 10-hydroxy-2E-decenoic acid: Possible epigenetic mechanisms of queen-worker bee differentiation. PLOS ONE 13, e0204538.

103. Alhosin, M. (2023). Epigenetics Mechanisms of Honeybees: Secrets of Royal Jelly. Epigenetics Insights 16, 25168657231213717.

104. Tschinkel, W.R. (1991). Insect sociometry, a field in search of data. Insectes Sociaux 38, 77–82.

105. Tschinkel, W.R. (1993). Sociometry and sociogenesis of colonies of the fire ant Solenopsis invicta during one annual cycle: ecological archives M063-002. Ecol. Monogr. 63, 425–457.

106. Bhatkar, A., and Whitcomb, W. (1970). Artificial diet for rearing various species of ants. Fla. Entomol., 229–232.

107. Bustin, S.A., Benes, V., Garson, J.A., Hellemans, J., Huggett, J., Kubista, M., Mueller, R., Nolan, T., Pfaffl, M.W., and Shipley, G.L. (2009). The MIQE Guidelines: Minimum Information for Publication of Quantitative Real-Time PCR Experiments.

108. Taylor, S., Wakem, M., Dijkman, G., Alsarraj, M., and Nguyen, M. (2010). A practical approach to RT-qPCR—publishing data that conform to the MIQE guidelines. Methods 50, S1–S5.

109. Rodríguez, A., Rodríguez, M., Córdoba, J.J., and Andrade, M.J. (2015). Design of primers and probes for quantitative real-time PCR methods. In PCR primer design (Springer), pp. 31–56.

110. Thornton, B., and Basu, C. (2015). Rapid and simple method of qPCR primer design. In PCR primer design (Springer), pp. 173–179.

111. Bustin, S., and Huggett, J. (2017). qPCR primer design revisited. Biomol. Detect. Quantif. 14, 19–28.

112. Ye, J., Coulouris, G., Zaretskaya, I., Cutcutache, I., Rozen, S., and Madden, T.L. (2012). Primer-BLAST: a tool to design target-specific primers for polymerase chain reaction. BMC Bioinformatics 13, 134.

113. Ponton, F., Chapuis, M.-P., Pernice, M., Sword, G.A., and Simpson, S.J. (2011). Evaluation of potential reference genes for reverse transcription-qPCR studies of physiological responses in Drosophila melanogaster. J. Insect Physiol. 57, 840–850.

114. Freitas, F.C., Depintor, T.S., Agostini, L.T., Luna-Lucena, D., Nunes, F.M., Bitondi, M.M., Simões, Z.L., and Lourenço, A.P. (2019). Evaluation of reference genes for gene expression analysis by real-time quantitative PCR (qPCR) in three stingless bee species (Hymenoptera: Apidae: Meliponini). Sci. Rep. 9, 17692.

115. Livak, K.J., and Schmittgen, T.D. (2001). Analysis of relative gene expression data using real-time quantitative PCR and the 2− ΔΔCT method. methods 25, 402–408.

116. Cassill, D.L., and Tschinkel, W.R. (1995). Allocation of liquid food to larvae via trophallaxis in colonies of the fire ant, Solenopsis invicta. Anim. Behav. 50, 801–813.

117. Bear, A., Prudic, K.L., and Monteiro, A. (2017). Steroid hormone signaling during development has a latent effect on adult male sexual behavior in the butterfly Bicyclus anynana. PLoS One 12, e0174403.

118. Black, K., Petruk, S., Fenstermaker, T.K., Hodgson, J.W., Caplan, J.L., Brock, H.W., and Mazo, A. (2016). Chromatin proteins and RNA are associated with DNA during all phases of mitosis. Cell Discov. 2, 16038.

